# Seclidemstat blocks the transcriptional function of multiple FET-fusion oncoproteins

**DOI:** 10.1101/2024.05.19.594897

**Authors:** Galen C. Rask, Cenny Taslim, Ariunaa Bayanjargal, Matthew V. Cannon, Julia Selich-Anderson, Jesse C. Crow, Aundrietta Duncan, Emily R. Theisen

## Abstract

Genes encoding the RNA-binding proteins <u>F</u>US, <u>E</u>WSR1, and <u>T</u>AF15 (FET proteins) are involved in chromosomal translocations in rare sarcomas. FET-rearranged sarcomas are often aggressive malignancies affecting patients of all ages. New therapies are needed. These translocations fuse the 5’ portion of the FET gene with a 3’ partner gene encoding a transcription factor (TF). The resulting fusion proteins are oncogenic TFs with a FET protein low complexity domain (LCD) and a DNA binding domain. FET fusion proteins have proven stubbornly difficult to target directly and promising strategies target critical co-regulators. One candidate is lysine specific demethylase 1 (LSD1). LSD1 is recruited by multiple FET fusions, including EWSR1::FLI1. LSD1 promotes EWSR1::FLI1 activity and treatment with the noncompetitive inhibitor SP-2509 blocks EWSR1::FLI1 transcriptional function. A similar molecule, seclidemstat (SP-2577), is currently in clinical trials for FET-rearranged sarcomas (NCT03600649). However, whether seclidemstat has pharmacological activity against FET fusions has not been demonstrated. Here, we evaluate the *in vitro* potency of seclidemstat against multiple FET-rearranged sarcoma cell lines, including Ewing sarcoma, desmoplastic small round cell tumor, clear cell sarcoma, and myxoid liposarcoma. We also define the transcriptomic effects of seclidemstat treatment and evaluated the activity of seclidemstat against FET fusion transcriptional regulation. Seclidemstat showed potent activity in cell viability assays across FET-rearranged sarcomas and disrupted the transcriptional function of all tested fusions. Though epigenetic and targeted inhibitors are unlikely to be effective as a single agents in the clinic, these data suggest seclidemstat remains a promising new treatment strategy for patients with FET-rearranged sarcomas.

**SIGNIFICANCE:** Here, we show the noncompetitive inhibitor, seclidemstat, has *in vitro* activity against multiple FET fusion proteins that cause a number of rare and aggressive sarcomas. These data represent one of the largest analyses of FET fusion activity across multiple malignancies and are a valuable resource for those studying FET-rearranged sarcomas.

## INTRODUCTION

Genes encoding the RNA-binding proteins <u>F</u>US, <u>E</u>WSR1, and <u>T</u>AF15 (FET family proteins) are frequently involved in chromosomal translocations in sarcomas.^1^ The resulting fusion genes typically involve a FET family gene in the 5’-portion of the fusion and a gene encoding a transcription factor (TF) in the 3’-portion of the fusion.^1,2^ The proteins expressed from these fusions include the amino-terminal intrinsically disordered and low complexity domain (LCD) of the FET protein and a carboxy-terminal DNA binding domain (DBD).^1,2^ The LCD functions as a strong transcriptional activator and these chimeras function as oncogenic transcription factors. Sarcomas with FET-rearrangements are thus often characterized by a relatively low mutational burden and widespread deregulation of chromatin and transcription.^2–11^

The epidemiology of tumors involving FET rearrangements varies. For example Ewing sarcoma, involving FET::ETS fusions (primarily EWSR1::FLI1), is predominantly a tumor of adolescents and young adults.^12^ This is an aggressive malignancy, and though patients with local disease have ∼70% 5-year survival, patients with metastatic, relapsed and refractory disease have dismal outcomes with 10-30% survival.^13,14^ Desmoplastic small round cell tumor (DSRCT) is a rare and aggressive mesenchymal tumor characterized by the expression of the EWSR1::WT1 fusion.^15^ DSRCT primarily affects adolescents and young adults, and has extremely poor outcomes with ∼60-70% of patients succumbing to disease within 3 years of diagnosis.^15^ Clear cell sarcoma is characterized by expression of EWSR1::ATF1 fusion protein and has a peak incidence in middle age.^16^ Despite multimodal therapy, 5-year survival is ∼50% and 10-year survival is ∼38%.^16^ Myxoid liposarcomas represent ∼30% of all liposarcomas and are characterized by the expression of the FUS::DDIT3 fusion.^17^ Peak incidence for this disease is in adults near 50 years old and 5-year survival is ∼75-85%.^17^ Outcomes are worse for those with metastatic or high-grade disease.^17^ Outcomes for patients with metastatic and relapsed FET-rearranged sarcomas have remained unchanged for decades and new therapeutics options are urgently needed.

As the causative oncogenes, these fusion proteins make attractive therapeutic targets. Unfortunately, the intrinsically disordered nature of the LCD and the convex binding surface of TF DBDs makes pharmacologically targeting these proteins difficult.^12^ Instead, inhibiting the critical regulators and co-regulators of the fusion protein have emerged as attractive therapeutic strategies.^18–23^ Toward this end, many chromatin co-regulators have been identified that interact with the N-terminal FET domain, including the ATP-dependent chromatin remodeling BAF complex^2,7,10,24^, the histone acetyltransferase p300^9^, histone deacetylases (HDACs)^25^, and the histone lysine demethylase LSD1^18,24,26^. Our prior results showed that treatment with the noncompetitive LSD1 inhibitor, SP-2509, reverses the transcriptional activity of both EWSR1::FLI1 and EWSR1::ERG in Ewing sarcoma.^18^ Notably, while treatment with SP-2509 blocked both fusion-dependent gene activation and repression, HDAC inhibition only blocked EWSR1::FLI1-mediated repression, highlighting the importance of finding the right epigenetic targeted therapy.^18^

Combined with results showing single agent activity in xenograft models^18^, these data spurred the development of a clinical lead compound with improved physicochemical and pharmacokinetic properties, seclidemstat (SP-2577). Seclidemstat is currently in clinical trials for Ewing sarcoma and related FET-rearranged tumors (NCT03600649, NCT05266196), as well as myelodysplastic syndrome (MDS) and chronic myelomonocytic leukemia (CMML) (NCT04734990). Though seclidemstat is structurally similar to SP-2509, its effects on the transcriptional activity of EWSR1::FLI1 have not been directly assessed and it is unknown whether its pharmacodynamic effects are comparable to SP-2509. Additionally, given reported similarities in the fusion-driven transcriptional changes in different FET-rearranged tumors^11^ and the demonstrated recruitment of LSD1 by multiple FET fusion proteins^24,26^ treatment with seclidemstat may also block the transcriptional activity of other fusions. We therefore tested whether treatment with the noncompetitive inhibitor seclidemstat blocks the transcriptional functions of varied FET fusion oncoproteins in *in vitro* models of multiple malignancies: Ewing sarcoma, DSRCT, clear cell sarcoma, and myxoid liposarcoma. These studies revealed that all disease models were sensitive to treatment with noncompetitive LSD1 inhibitors and that seclidemstat blocked FET fusion activity in all the cell lines tested, as seen for SP-2509 in Ewing sarcoma.

## MATERIALS AND METHODS

### Reagents and constructs

SP-2509 and OG-L002 were purchased from Selleck Chemicals (Cat. # S7680 and S7237, respectively.) Seclidemstat was provided by Salarius Pharmaceuticals. SP-2513 was synthesized by the Medicinal Chemistry Shared Resource at The Ohio State University James Comprehensive Cancer Center. Identity and purity of SP-2513 were verified by HPLC-MS and NMR. The luciferase RNAi (iLuc), pMSCV-empty vector (hygro), and pMSCV-3X-FLAG EWSR1::FLI1 (hygro) are previously described.^25,33–39^ The EWSR1::ERG RNAi construct is previously described^34^, as is the pMSCV 3X-FLAG EWSR1::ERG construct.^25^

### Cell lines and tissue culture

Ewing sarcoma cell lines A673, TTC-466, SK-N-MC and TC32 were provided by Dr. Stephen Lessnick. The DSRCT cell lines JN-DSRCT-1 and BER were kindly provided by Dr. Sean B. Lee. The myxoid liposarcoma cell lines 1765-92 and 402-91, as well as the clear cell sarcoma cell line DTC1 were kindly provided by Dr. Torsten Nielsen. The clear cell sarcoma cell line SU-CCS-1 was purchased from ATCC (#CRL-2971). DL221 was provided by Salarius Pharmaceuticals. All cells were maintained at 37°C, 5% CO2. A673 cells were cultured in Dulbecco’s Modified Eagle Medium (DMEM; Corning Cellgro 10-013-CV) supplemented with 10% fetal bovine serum (FBS; Gibco 16000-044), penicillin/streptomycin/glutamine (P/S/Q; Gibco 10378-016), and sodium pyruvate (Gibco 11360-070). DL221 cells were cultured in DMEM supplemented with 10% FBS and PSQ. JN-DSRCT-1 and BER cells were grown in DMEM/F12 media with 10% FBS and PSQ. All of the remaining cell lines (TTC-466, 1765-92, 402-91, SU-CCS-1, and DTC1) were cultured in RPMI with 10% FBS and PSQ. All experiments were performed within 2 months of thawing a cell line. All cell lines were tested for mycoplasma using the Universal Mycoplasma Detection Kit (ATCC 30-1012K) and STR profiled (LabCorp) upon thawing and are checked annually if in regular use in the lab.

### Retrovirus production and transduction of TTC-466 cells

Viral production using HEK-293 cells was performed as previously described.^18,25,33,37,39^ shRNAs and cDNAs were transduced on day 1, with cDNA transduction beginning 4 hours following shRNA transduction. Transduction was performed with viral supernatants containing 2.5 µL of 8 µg/mL polybrene. Plain media was added overnight and on day 3 cells were passaged into selection media containing 2 µg/uL puromycin (Sigma P8833) and 150 µg/mL hygromycin B (Thermo 10687010) for 10 days. Following 10 days, cells were seeded into soft agar assays and collected for RNA and protein isolation.

### Antibodies

The following antibodies were used for immunodetection: M2-anti-FLAG (Sigma F3165), anti-ERG (Abcam ab92513), anti-α-tubulin (DM1A; Abcam ab7291), IRDye® 800CW goat anti-mouse IgG (LI-COR Biosciences 926-32210), and IRDye® 800CW goat anti-rabbit IgG (LI-COR Biosciences 926-32211).

### Cell viability assays

Cell viability was assessed by seeding cells in 96-well white, opaque, tissue-culture treated plate (Corning) at a density of 10,000 cells per well in 200µl of corresponding media. After 24 hours on the plate, cells were treated independently with a 9-point 3.3x increasing dosing scheme of SP-2509, seclidemstat, SP-2513 and OG-L002 (2nM-30µM). After 96h, 122µl of media was removed from each well and 80µl of CellTiter-Glo (Promega) was added to each well. Plates were agitated while covered at 250rpm for 10 minutes at room temperature. Luminescence was measured on a Synergy H1 plate reader (Agilent). Cell viability was calculated relative to vehicle treated wells.

### Protein isolation and western blot

Total cellular protein was extracted from frozen cell pellets using the Pierce™ RIPA buffer (ThermoFisher 88901) with protease inhibitor cocktail added (Sigma P8340). Pellets were resuspended in RIPA for 30 minutes, vortexed vigorously every 10 minutes, and centrifuged at max speed for 30 minutes at 4°C. Protein concentration was determined with the Pierce™ BCA Protein Assay Kit (Thermo Scientific 23225). 15 µg of protein sample was run on precast 4-15% gradient Tris-Glycine gels (BioRad 4561084) at 90V for 15 minutes and 120V for 60 minutes. Proteins were transferred to nitrocellulose membranes (Thermo IB23002) using the iBlot™ 2 (Thermo IB21001). Membranes were blocked with Odyssey® PBS Blocking Buffer (Li-Cor 927-40003) for 1 hour at room temperature. Immunoblotting was performed with primary antibody overnight at 4°C.

### Soft Agar Assays

Soft agar assays were performed as previously described^25,29,37^, with 18,000 cells/plate seeded for TTC-466 cells.

### RNA-isolation and RNA-seq submission

Cells were seeded with 1×10^6^ cells in 10 mL media per 10-cm dish for RNA-seq experiments. After 24 hours, media was replaced with fresh media containing either vehicle (0.3% DMSO) or the IC90 of seclidemstat as outlined in Table 1. Cells were treated for 48 hours, trypsinized, washed in 1X Hank’s Balanced Salt Solution (HBSS), pelleted and flash frozen for RNA extraction.

**Table 1.** FET-rearranged cell line models and RNA-seq dosing used for this study.

| Tumor | Cell line | Fusion | Seclidemstat<br>Dose for RNA-seq (nM) |
| --- | --- | --- | --- |
| Ewing Sarcoma | A673 | EWSR1::FLI1 | 1076 |
|  | TC32 | EWSR1::FLI1 | N/A |
|  | SK-N-MC | EWSR1::FLI1 | N/A |
|  | TTC-466 | EWSR1::ERG | 1069 |
| DSRCT | JN-DSRCT-1 | EWSR1::WT1 | 1265 |
|  | BER | EWSR1::WT1 | 2546 |
| Clear Cell Sarcoma | SU-CCS-1 | EWSR1::ATF1 | 802.2 |
|  | DTC1 | EWSR1::ATF1 | 2758 |
| Myxoid<br>Liposarcoma | 1765-92 | FUS::DDIT3 | 2766 |
|  | 402-91 | FUS::DDIT3 | 1452 |
|  | DL221 | FUS::DDIT3 | 14091 |

Total mRNA was extracted from cell pellets following the manufacturer’s instructions of the RNeasy kit (Qiagen) including the use of a gDNA removal column. RNA concentration was quantified using a Nanodrop and 1 µg of RNA per sample was submitted to the Abigail Wexner Research Institute Genomic Services Laboratory for library preparation using the TruSeq Stranded mRNA Kit (Illumina 20020594), library quality control, and next generation sequencing using an Illumina NovaSeq 6000. Samples were sequenced with a targeted sequencing depth of 50 million 150-bp paired end reads per sample. Raw BCL files were converted to FASTQ files and returned following the sequencing runs.

### RNA-seq data analysis

Publicly available data from FASTQ files were analyzed using an in-house containerized pipeline that performs quality control (FastQC^63^, MultiQC^64^), adapter and low quality read trimming (TrimGalore!^65^), alignment to hg38 reference genome (STAR 2.6^66^) and assignment to genomic features (featureCounts^67^), and analyzed for differential expression (DESeq2^68^, SARTools^69^). Fusion detection was run using EnFusion as described.^31^ Overlap analysis was done using GeneOverlap^70^ and visualized using eulerr^71^, venn^72^, and UpSetR^73^. UMAP was plotted using the R umap package 16.^74^ Other visualization was done with ggplot2^75^ and base R packages. Pathway analysis was performed using clusterProfiler^76^ and misgdb database^77^.

### Statistical Analysis

Dose response curves for cell viability data were determined in GraphPad Prism 9 using the 4-parameter, variable slope log(inhibitor) vs. response equation. Three technical replicates were included in each of two biological replicates. Differentially expressed genes were defined as genes with a multiple hypothesis testing adjusted (FDR/Benjamini-Hochberg^78^) p-value of <0.05. Statistical significance of Venn overlap analysis was evaluated using the Jaccard index^79,80^, which measures the similarity and Fisher’s exact test for statistical significance.^81^ GSEA was performed as described^32^, with normalized enrichment scores and p-values reported.

### Data availability

The data sets analyzed in this work are deposited in the NCBI’s Gene Expression Omnibus and Sequencing Read Archive and are accessible through GEO Series Accession GSE267611.

## RESULTS

### Noncompetitive LSD1 inhibitors show potent activity against cell lines derived from FET-rearranged tumors

In order to determine the effects of seclidemstat on the transcriptional activity of different FET fusion proteins, we first sought to define the potency of seclidemstat against cell lines expressing these fusions in cell viability assays. We used the following cell lines: Ewing sarcoma – A673, TC32, SK-N-MC, TTC-466; DSRCT – JN-DSRCT-1, BER; clear cell sarcoma – SU-CCS-1, DTC1; and myxoid liposarcoma – 1765-92, 402-91, DL221. The respective fusions expressed in each cell line are shown in Table 1. In addition to seclidemstat, we included SP-2509 for comparison to another noncompetitive LSD1 inhibitor as well as OG-L002 for comparison to an irreversible LSD1 inhibitor with similar potency and specificity^27^. Lastly, we included a compound structurally similar to SP-2509 with no activity against LSD1, SP-2513 (cpd 13^28^), to exclude off-target cytotoxic effects caused by the hydrazide moiety in seclidemstat and SP-2509.

As previously reported^29^, all Ewing sarcoma cell lines were sensitive to treatment with SP-2509, with IC50s ranging from 30-500 nM, and resistant to treatment with OG-L002 with no determinable IC50 up to 30 µM (Figure 1A-B, Supplementary Figure 1). Ewing sarcoma cells were also sensitive to seclidemstat with IC50s ranging from 290-700 nM (Figure 1A-B, Supplementary Figure 1). This slight decrease in *in vitro* potency is expected. The structure of SP-2509 contains a morpholine ring, which promotes cell permeability, while seclidemstat instead possesses an N-methylpiperazine.^28,30^ This moiety improves formulation, solubility, oral bioavailability, and pharmacokinetics, but can reduce permeability in tissue culture assays. Notably, SP-2513 showed no activity in cell viability assays, with IC50s not determinable below 30 µM (Figure 1A-B, Supplementary Figure 1).

**Figure 1.**
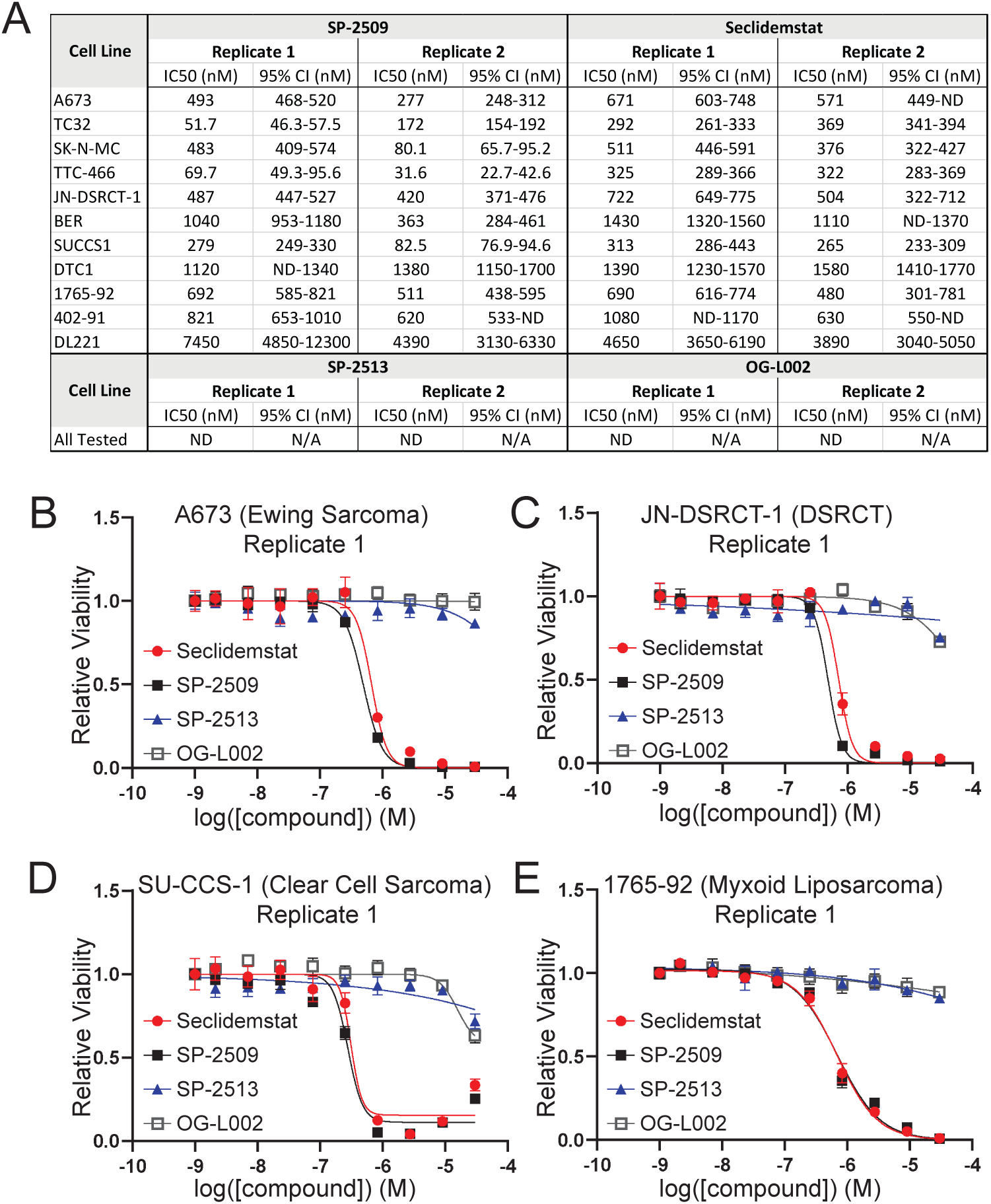
Noncompetitive LSD1 inhibitors show potent activity in cell viability assays using FET-rearranged sarcoma cell lines. (A) Compiled IC50s for SP-2509, seclidemstat, SP-2513, and OG-L002 for all tested cell lines and replicates. IC50s were determined using GraphPad Prism 9 and are reported with the 95% confidence interval. (B-E) Dose response curves for seclidemstat (red/circle), SP-2509 (black/closed square), SP-2513 (blue/triangle), and OG-L002 (gray/open square) in (B) A673, (C) JN-DSRCT-1, (D) SU-CCS-1, and (E) 1765-92 cells. Data are displayed for a single biological replicate. Mean values of 3 technical replicates are shown with standard deviation. Calculated curves of best fit are also shown.

A similar pattern of activity emerged in DSRCT and clear cell sarcoma. IC50s for SP-2509 and seclidemstat were 300 nM-1.1 µM and 500 nM-1.5 µM, respectively, in both DSRCT cell lines tested (Figure 1A, 1C, Supplementary Figure 2A-C). Clear cell sarcoma cell lines had IC50s ranging from 80 nM-1.4 µM and 250 nM-1.6 µM for SP-2509 and seclidemstat, respectively (Figure 1A, 1D, Supplementary Figure 2D-F). IC50s were undeterminable for OG-L002 and SP-2513 in these cell lines below 30 µM (Figure 1A, 1C-D, Supplementary Figure 2A-F). Similarly, neither OG-L002 nor SP-2513 showed activity in any of the myxoid liposarcoma cell lines (Figure 1A, 1E, Supplementary Figure 2G-K). There was more variable potency observed for SP-2509 and seclidemstat in the myxoid sarcoma cell lines. 1765-92 and 402-91 had IC50s ranging from 500-850 nM for SP-2509 and 450 nM-1.1 µM for seclidemstat. In contrast, the DL221 cell line was more resistant. We were unable to fit a 4-parameter dose response curve, because there was not enough data to establish the bottom of the curve below 30 µM, above which we encounter challenges related to solubility (Supplementary Figure 2J-K). Instead, we fixed the bottom of the curve to 0 and used the estimated IC50. The IC50s across replicates ranged from 4.2-7.5 µM for SP-2509 and 3.8-4.7 µM for seclidemstat.

Taken together, these data show that seclidemstat has comparable potency to SP-2509 in all the cell lines tested. Moreover, other FET-rearranged tumors showed a similar pattern of activity, with sensitivity to the noncompetitive LSD1 inhibitors, but no sensitivity to the irreversible or catalytic inhibitor OG-L002. This suggest that cytotoxic activity does not require inhibition of LSD1 enzymatic activity. The exception was the DL221 myxoid liposarcoma cell line, which was somewhat resistant to noncompetitive inhibitors. Nonetheless, SP-2509 and seclidemstat showed more activity than OG-L002 and SP-2513 in this cell line.

### Additional bioinformatic tools confirm cell line and data integrity in transcriptomic analyses

We used our calculated dose response curves to estimate an IC90 for each cell line and used this dose for our transcriptomic analyses (Table 1) to match what had been done previously with SP-2509 in Ewing sarcoma cells.^18^ Cells were treated for 48 hours with either the DMSO vehicle or seclidemstat prior to harvest for RNA-seq analysis. Though we regularly perform STR profiling on all the cell lines used in the lab, some cell lines here (DTC1 and DL221) did not have reference profile publicly available. Therefore, as another layer of quality control, we used the EnFusion pipeline to detect fusion transcripts expressed in each cell line from standard RNA seq data.^31^ This pipeline confirmed expression of *EWSR1::FLI1* and *EWSR1::ERG* in A673 and TTC-466 cell lines, respectively (Supplementary Figure 3A-B). *EWSR1::WT1* was detected in both JN-DSRCT-1 and BER with different breakpoints predicted for each cell line (Supplementary Figure 4A-B). Both SU-CCS-1 and DTC1 were confirmed to express *EWSR1::AFT1* with the same predicted breakpoint (Supplementary Figure 5A-B). Notably, despite the large differences seen between DL221 and the other two mxyoid liposarcoma cell lines, all three had detectable expression of *FUS::DDIT3* (Supplementary Figure 6A-C). As a final step of quality control and to ensure proper handling of samples and data, we used uniform manifold approximation and projection (UMAP) to assess all our RNA-seq samples used for analysis. We found that samples, either treated or untreated, clustered together first by disease and second by cell line (Supplementary Figure 7). These data indicate that inter-disease variability drove the largest differences in transcriptomes across samples, as expected.

### Seclidemstat treatment alters transcription in a similar manner as SP-2509 in Ewing sarcoma

Having confirmed appropriate fusion expression and sample clustering, we analyzed our differential expression data for cells treated with seclidemstat. We first asked whether treatment with seclidemstat was similar to treatment with SP-2509. To do this, we reanalyzed our most recent SP-2509 data from Pishas, *et. al.*, 2018^29^ with our current RNA-seq pipeline for comparison to seclidemstat. We found that both SP-2509 and seclidemstat upregulate and downregulate the expression of thousands of genes with significant overlap in the differentially expressed genes (DEGs) in both treatment conditions (Supplementary Figure 8A-C). Likewise, using gene set enrichment analysis (GSEA^32^) we found significant functional similarity in seclidemstat and SP-2509 downregulated genes (normalized enrichment score [NES]: -3.0488, Supplementary Figure 8D) and upregulated genes (NES: 2.7995, Supplementary Figure 8E). These data demonstrate significant similarity in the transcriptional changes caused by treatment with both seclidemstat and SP-2509 in A673 Ewing sarcoma cells.

Having shown that seclidemstat largely recapitulates the transcriptional effects of SP-2509, we also wanted to determine the degree to which seclidemstat disrupts EWSR1::FLI1 transcriptional activity and LSD1 function in Ewing sarcoma cells. As for SP-2509 above, we reprocessed previously published data for EWSR1::FLI1- and LSD1-regulated genes^29,33^ with our current pipeline. Importantly, this analysis of EWSR1::FLI1 gene regulation uses only DEGs detected upon rescue with EWSR1::FLI1 in a knockdown (KD)-rescue experiment. This restricts analysis to genes regulated by EWSR1::FLI1 expression and excludes genes whose expression changes as an off target of RNAi-mediated knockdown. Four-way Venn overlaps for SP-2509, seclidemstat, EWSR1::FLI1, and LSD1 DEGs are shown in Supplementary Figure 9A-B. Analysis of these data show significant overlap in SP-2509 and seclidemstat treatment as seen above, as well as significant overlaps in genes downregulated by seclidemstat and upregulated by LSD1 and vice versa (Supplementary Figure 9C). Interestingly, there is also some overlap between seclidemstat and LSD1 downregulated genes, as well as seclidemstat and LSD1 upregulated genes, a pattern not seen with SP-2509 (Supplementary Figure 9C). The association between seclidemstat treatment and reversal of LSD1 function is stronger (Supplementary Figure 9C). This is confirmed using GSEA, where LSD1-upregulated genes are downregulated by seclidemstat (NES: -2.4373; Supplementary Figure 9D) and LSD1-downregulated genes are upregulated by seclidemstat (NES: 2.16; Supplementary Figure 9E).

As for LSD1 the effects of seclidemstat treatment on EWSR1::FLI1 transcriptional regulation showed significant impact of seclidemstat upregulation on both EWSR1::FLI1-activated and -repressed genes (Supplementary Figure 9C). The same was true for seclidemstat downregulation (Supplementary Figure 9C). Interestingly and in contrast to LSD1 above, both seclidemstat up- and downregulation were more strongly overlapped with EWSR1::FLI1-activated genes than repressed genes. This finding was similar to our GSEA results, with EWSR1::FLI1-activated and repressed genes both more strongly associated with seclidemstat upregulation (NES: 1.8047 and NES: 1.3087, respectively; Supplementary Figure 9F-G). There are several factors that may contribute to these findings. First is the reduced cell permeability expected with seclidemstat as compared to SP-2509, and second is the relatively lower dose used for seclidemstat (∼1 µM) as compared to SP-2509 (2 µM^29^). Both may reduce the effective dose of seclidemstat in the same tissue culture conditions and this is supported by the overall lower number of seclidemstat-regulated genes as compared to SP-2509 (Supplementary Figure 8A-B).

Nonetheless, treatment with seclidemstat significantly impacts both EWSR1::FLI1 and LSD1-regulated genes, and closely resembles SP-2509 treatment and this is further validated at the pathway level. Pathway analysis using the MSigDB Curated Gene Sets shows that molecular signatures of EWSR1::FLI1-downregulation, mesenchymal tissue, and bone progenitors are all downregulated by EWSR1::FLI1 and LSD1 in our data, and upregulated by SP-2509 and seclidemstat (Supplementary Figure 10). Likewise, signatures related to proliferation and stem cell programs are downregulated by both SP-2509 and seclidemstat and upregulated by EWSR1::FLI1 and LSD1 (Supplementary Figure 10). Indeed, the pathways upregulated by EWSR1::FLI1 and LSD1 are largely downregulated by seclidemstat and vice versa (Supplementary Figure 10). These data show that seclidemstat treatment significantly alters gene regulation in Ewing sarcoma cells with a potency and in a manner similar to that reported for SP-2509.^18,29^

### Treatment with seclidemstat broadly alters gene regulation in FET-rearranged sarcomas

Given the activity observed in cell viability assays, we next asked whether seclidemstat also altered gene regulation in cell lines with other FET fusion proteins. As seen for A673 cells, seclidemstat treatment changed the expression of thousands of genes in all the tested cell lines (Figure 2A-L). This was true for both upregulated genes (Figure 2A, 2D, 2G, 2J) and downregulated genes (Figure 2B, 2E, 2H, and 2K). Additionally, there was significant functional overlap (i.e. common up DEGs and common down DEGs) in the genes regulated by seclidemstat across cell lines within each disease category (Figure 2C, 2F, 2I, 2L).

**Figure 2.**
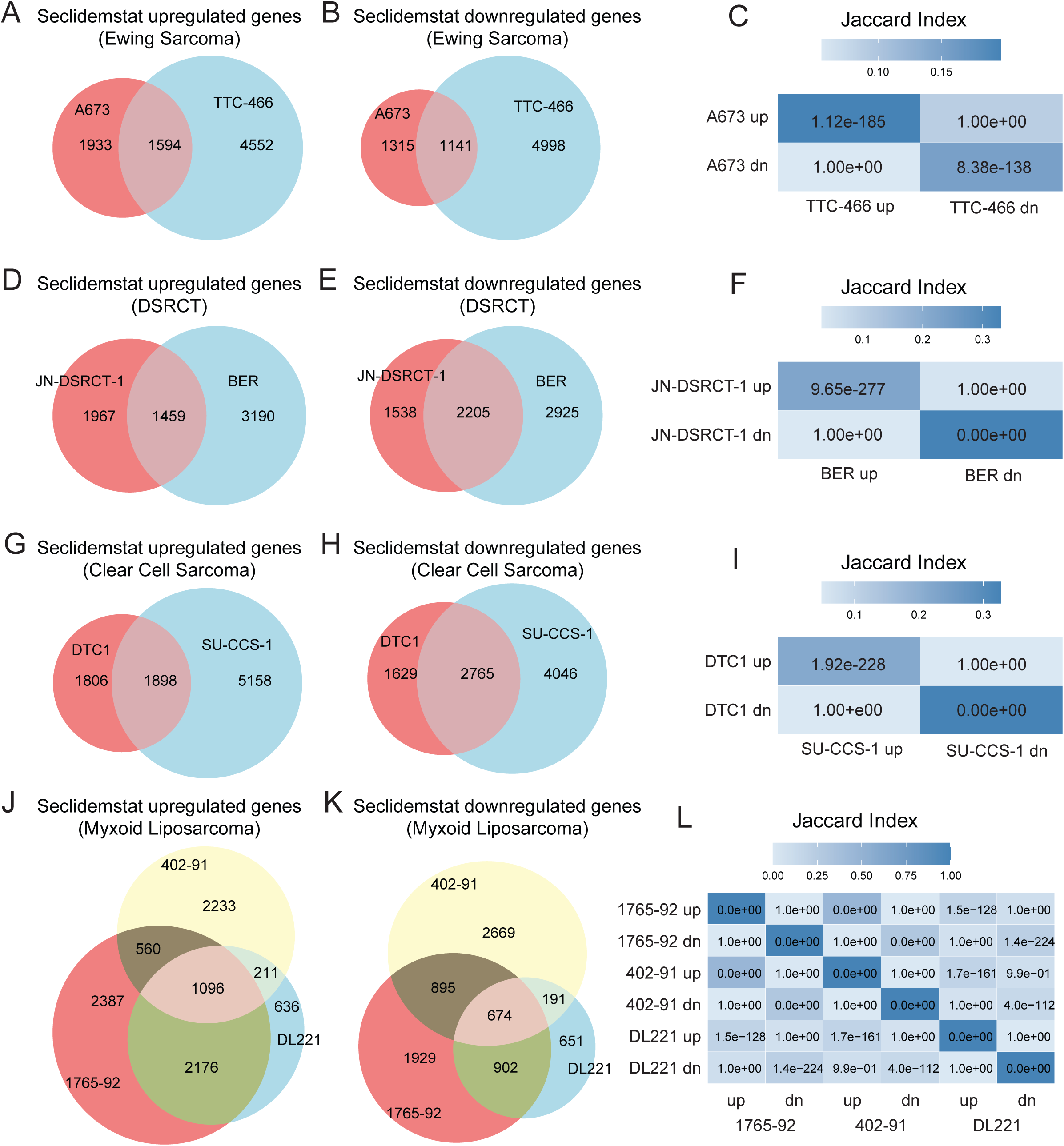
Seclidemstat alters the expression of thousands of genes in FET-rearranged sarcoma cell lines. (A-C) Venn overlap analysis of seclidemstat (A) up- and (B) downregulated genes in A673 and TTC-466 Ewing sarcoma cell lines with the Jaccard index and p-values of overlap (numbers inside each cell) shown in (C). (D-F) Venn overlap analysis of seclidemstat (D) up- and (E) downregulated genes in JN-DSRCT-1 and BER DSRCT cell lines with the Jaccard index and p-values of overlap (numbers inside each cell) shown in (F). (G-I) Venn overlap analysis of seclidemstat (G) up- and (H) downregulated genes in DTC1 and SU-CCS-1 clear cell sarcoma cell lines with the Jaccard index and p-values of overlap (numbers inside each cell) shown in (I). (J-L) Venn overlap analysis of seclidemstat (J) up- and (K) downregulated genes in 1765-92, 402-91, and DL221 myxoid liposarcoma cell lines with the Jaccard index and p-values of overlap (numbers inside each cell) shown in (L).

Though there was significant similarity in genes regulated by seclidemstat within a given disease, we also wanted to understand the genes and pathways commonly regulated by seclidemstat across all tested cell lines. We first used an upset plot to look at commonly regulated genes (Figure 3A-B). These analyses revealed that, for both seclidemstat-upregulated and -downregulated genes, the predominant DEGs were unique to each cell line (Figure 3A-B). There were also DEGs that were unique to each disease type, but these were typically far less abundant than DEGs unique to each cell line (Figure 3A-B). There were so few commonly upregulated DEGs (n=45) that that category was not included in the top 40 categories in the upset plot (Figure 3A). For downregulated DEGs, we detected 98 common DEGs across all cell lines (Figure 3B).

**Figure 3.**
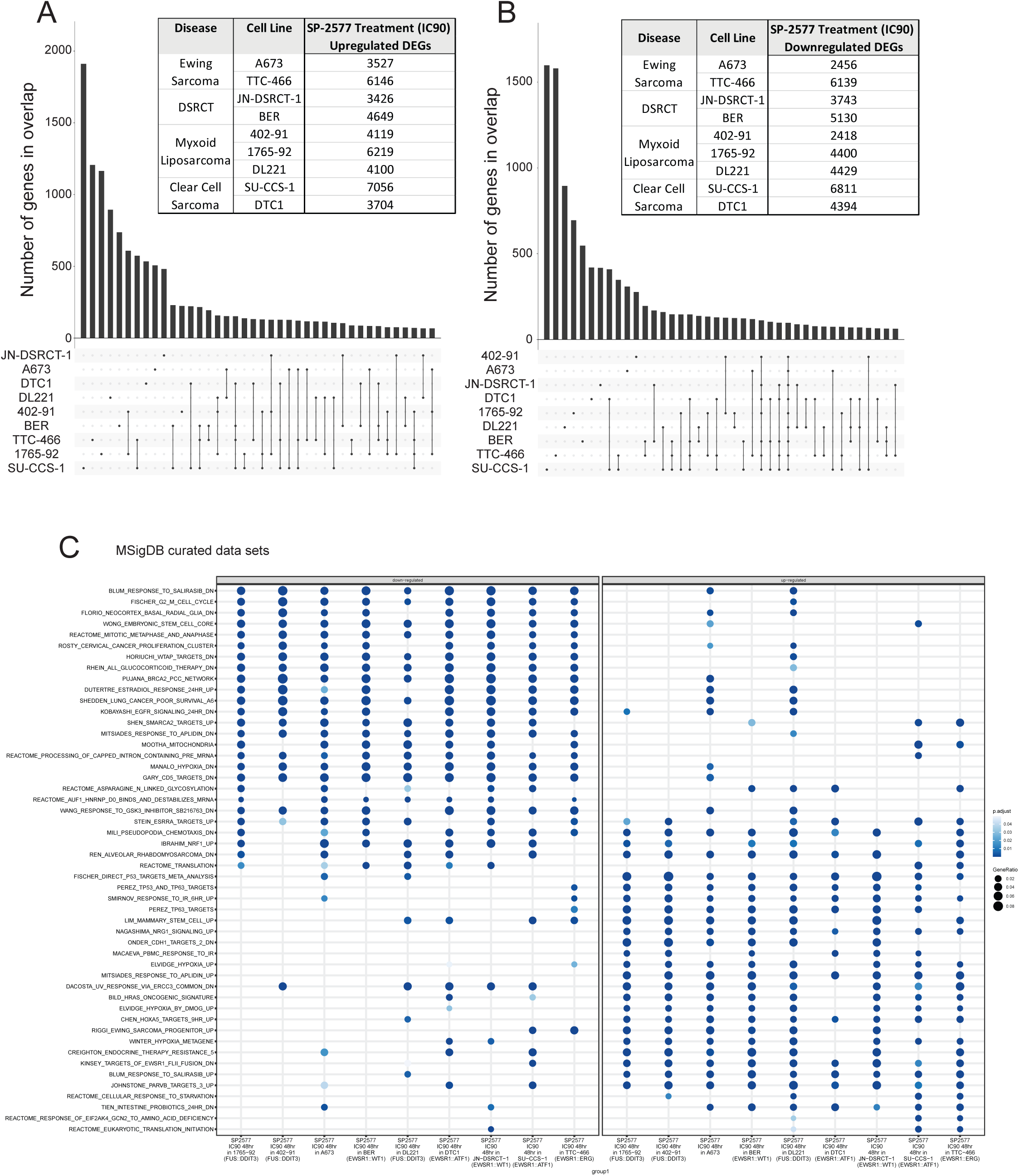
Seclidemstat regulates unique gene sets in each cell line and common biological pathways across cell lines. (A,B) Overlap analysis via upset plots of seclidemstat (A) up- and (B) downregulated genes in all 9 cell lines tested. Inset tables show the total number of differentially expressed genes detected in each cell line. (C) Pathway analysis for seclidemstat-regulated genes visualized with a dot plot using MSigDB curated gene signatures.

We next used pathway analysis to evaluate the relevant pathways regulated by seclidemstat treatment across cell lines. For this analysis we tested pathway signatures from MSigDB Curated Gene Sets (MSig; Figure 3C), Gene Ontology Biological Processes (GOBP; Supplementary Figure 11A), and Gene Ontology Molecular Function (GOMF; Supplementary Figure 11B). Gene signatures related to cell cycle, RNA processing, mitochondria and metabolism, embryonic stem cell programs, DNA replication, and transcription factor and co-factor binding were downregulated following treatment with seclidemstat (Figure 3, Supplementary Figure 11A-B). In contrast, gene signatures related to cell signaling, development, EWSR1::FLI1 and PAX3::FOXO1 repressed targets, and the cellular response to starvation were all upregulated in seclidemstat treated cells (Figure 3, Supplementary figure 11A-B). Considered together, these analyses suggest that, regardless of somewhat unique DEG sets in each cell line, seclidemstat treatment downregulates cellular pathways involved in oncogenesis, such as increased proliferation and a more dedifferentiated state, and upregulated pathways involved in cell signaling, differentiation, and development that are repressed by fusion oncogenes. These findings suggest that there may be common mechanisms of transcriptional regulation in FET-rearranged tumors that are disrupted by treatment with noncompetitive LSD1 inhibitors.

### FET fusion proteins show significant similarities in gene regulation

To evaluate the impact of seclidemstat treatment on the transcriptional function of FET fusion oncogenes, we first reanalyzed publicly available FET fusion transcriptional signatures for A673^33^ (as described above), JN-DSRCT-1^11^, BER^11^, and DTC1^6^. We also generated a new fusion regulated signature for EWSR1::ERG in TTC-466 cells using a similar approach as that described above for A673s. Briefly, we performed a standard KD-rescue experiment using a 3’UTR targeted shRNA to deplete endogenous EWSR1::ERG and rescued with a shRNA-resistant cDNA encoding EWSR1::ERG, similar to that reported for EWSR1::FLI1 in prior experiments.^33–39^ TTC-466 cells tolerated EWSR1::ERG KD/rescue with expression of the rescue construct at similar expression levels seen for both EWSR1::FLI1 and EWSR1:ERG (Supplementary Figure 12A), rescue of growth in soft agar comparable to that previously seen for EWSR1::FLI1 (Supplementary Figure 12B-C), and rescue of an oncogenic transcriptional signature (Supplementary Figure 12D). The EWSR1::ERG signature was then defined as genes differentially expressed in cells expressing the EWSR1:ERG rescue construct compared to the knockdown condition.

Overlap analyses of all five fusion transcriptional signatures revealed a large degree of overlap with 847 genes commonly upregulated and 351 genes commonly downregulated (Figure 4A-B) and statistically significant overlaps in each pairwise comparison between the respective up- and downregulated genes from each fusion (Figure 4C). As has been reported by Gedminas, *et. al.*^11^, there is significant overlap in the DEGs signatures for EWSR1::WT1 in both JN-DSRCT-1 and BER, as well as between both DSRCT cell lines and the EWSR1::FLI1 signature (Figure 4C). Also, as reported by Sankar, *et. al.*^18^, there is a significant overlap in the EWSR1::FLI1 and EWSR1::ERG transcriptional signatures (Figure 4C). Given these data, we were unsurprised to also see significant overlap between both EWSR1::WT1 DEG signatures and EWSR1::ERG DEGs (Figure 4C). Notably, the EWSR1::ATF1 DEG signature also showed significant overlap with all other fusions tested (Figure 4C). Despite these significant overlaps, the most prevalent DEG sets were unique to each cell line (Figure 4D-E). The next most prevalent DEG sets were those common to each disease and then those common to all fusions for both up- and downregulated genes (Figure 4D-E). There were more commonly upregulated genes than downregulated genes, and this may reflect common mechanisms of transcriptional activation in the presence of these fusions.

**Figure 4.**
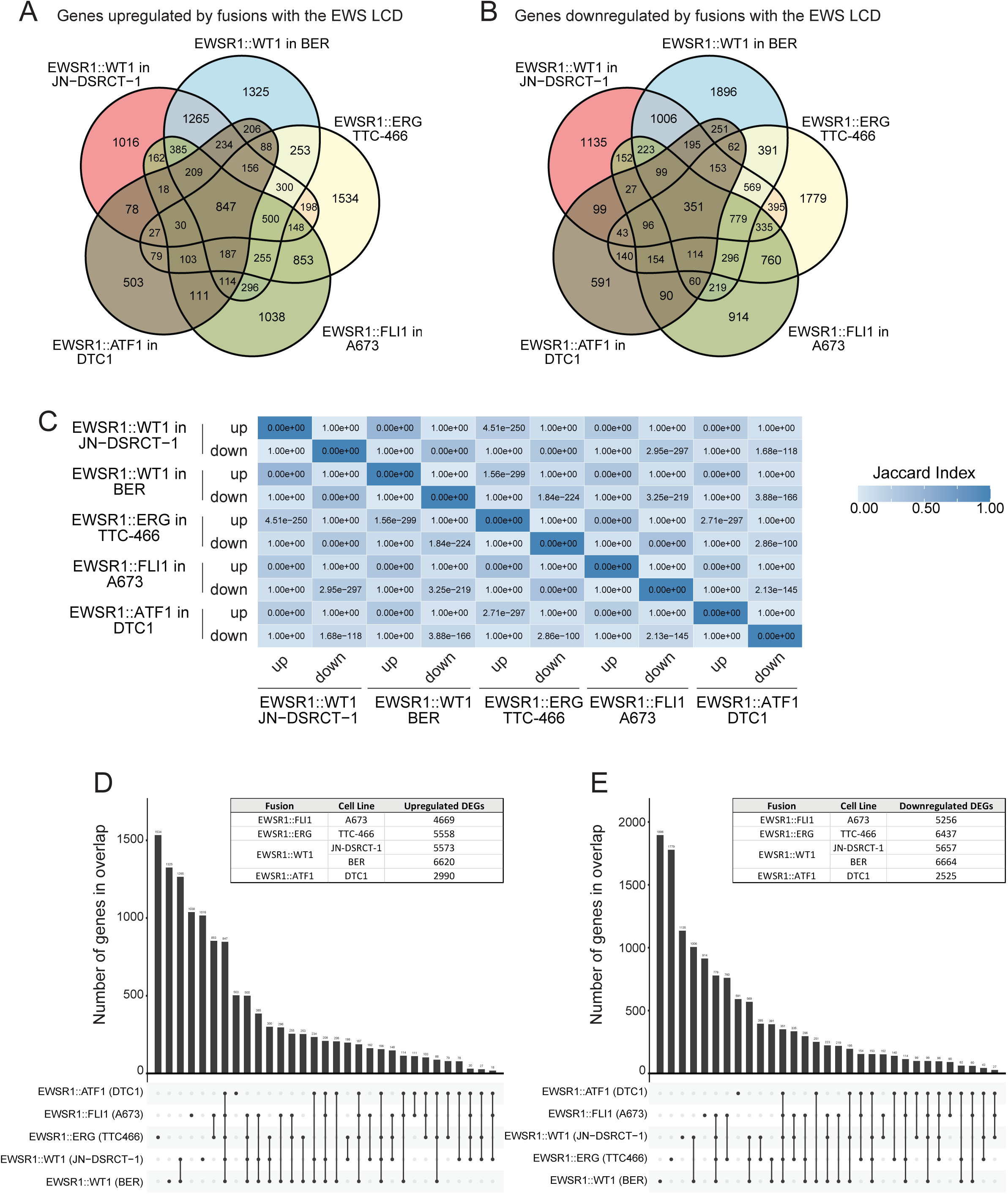
Different FET fusion proteins regulate common gene sets. (A-C) Venn overlap analysis of FET fusion (A) activated and (B) repressed genes in A673, TTC-466, JN-DSRCT-1, BER, and DTC1 cell lines with the Jaccard indices for pairwise comparisons and p-values of overlap (numbers inside each cell) shown in (C). (D,E) Overlap analysis via upset plots of FET fusion (D) activated and (E) repressed genes in all 5 cell lines tested. Inset tables show the total number of differentially expressed genes detected in each cell line.

Pathway analysis reinforced our findings of common gene regulation by all the fusions analyzed here. GOBP and GOMF showed that expression of FET fusion proteins resulted in the downregulation of genes related to development, autophagy, phospholipid binding, and extracellular matrix and adhesion (Supplementary Figure 13A-B). In contrast genes related to ribosome biogenesis and translation, RNA processing, DNA replication, and transcriptional coregulators were commonly upregulated with FET fusion expression. These findings are largely concordant with FET fusions disrupting the normal enhancer landscape that determines cell development and differentiation, high rates of disease dissemination, and widespread deregulation of transcription as has been reported.^6–10,15,16,40–44^

### Seclidemstat treatment reverses FET fusion transcriptional activity

In our last analysis, we asked whether seclidemstat treatment blocks the transcriptional activity of FET fusion proteins in other diseases. In both JN-DSRCT-1 and BER cells, we found that seclidemstat downregulated a significant number of EWSR1::WT1-activated genes and upregulated a significant number of EWSR1::WT1 downregulated genes (Figure 5A-C, Supplementary Figure 14A-C). Unlike the results for EWSR1::FLI1 discussed above, seclidemstat downregulation showed no significant overlap with EWSR1::WT1-repressed genes, and the same was true for seclidemstat upregulation of EWSR1::WT1-activated genes (Figure 5C, Supplementary Figure 14C). Heatmap analyses in Figure 5D and Supplementary Figure 14D, likewise show a reversal of the EWSR1::WT1 transcriptional signature following seclidemstat treatment. The results were largely the same for EWSR1::ATF-regulated genes in DTC1 cells (Figure 5E-H) and EWSR1::ERG-regulated genes in TTC-466 cells (Supplementary Figure 14E-H). In these cell lines, seclidemstat upregulated genes that were repressed by the fusion and vice versa, and there was no significant overlap between seclidemstat upregulation and fusion-activated genes or between seclidemstat downregulation and fusion-repressed genes. These data indicate that seclidemstat reverses the transcriptional activity of multiple FET fusion proteins, specifically EWSR1::FLI1, EWSR1::ERG, EWSR1::WT1, and EWSR1::ATF1. This finding is largely consistent with our observations in cell viability assays in Figure 1 and Supplementary Figures 1-2, the common set of pathways that change upon seclidemstat treatment as shown in Figure 3C and Supplementary Figure 11, and the degree of similarity in the genes that are regulated by the different fusions in different diseases as shown in Figure 4 and Supplementary Figure 13.

**Figure 5.**
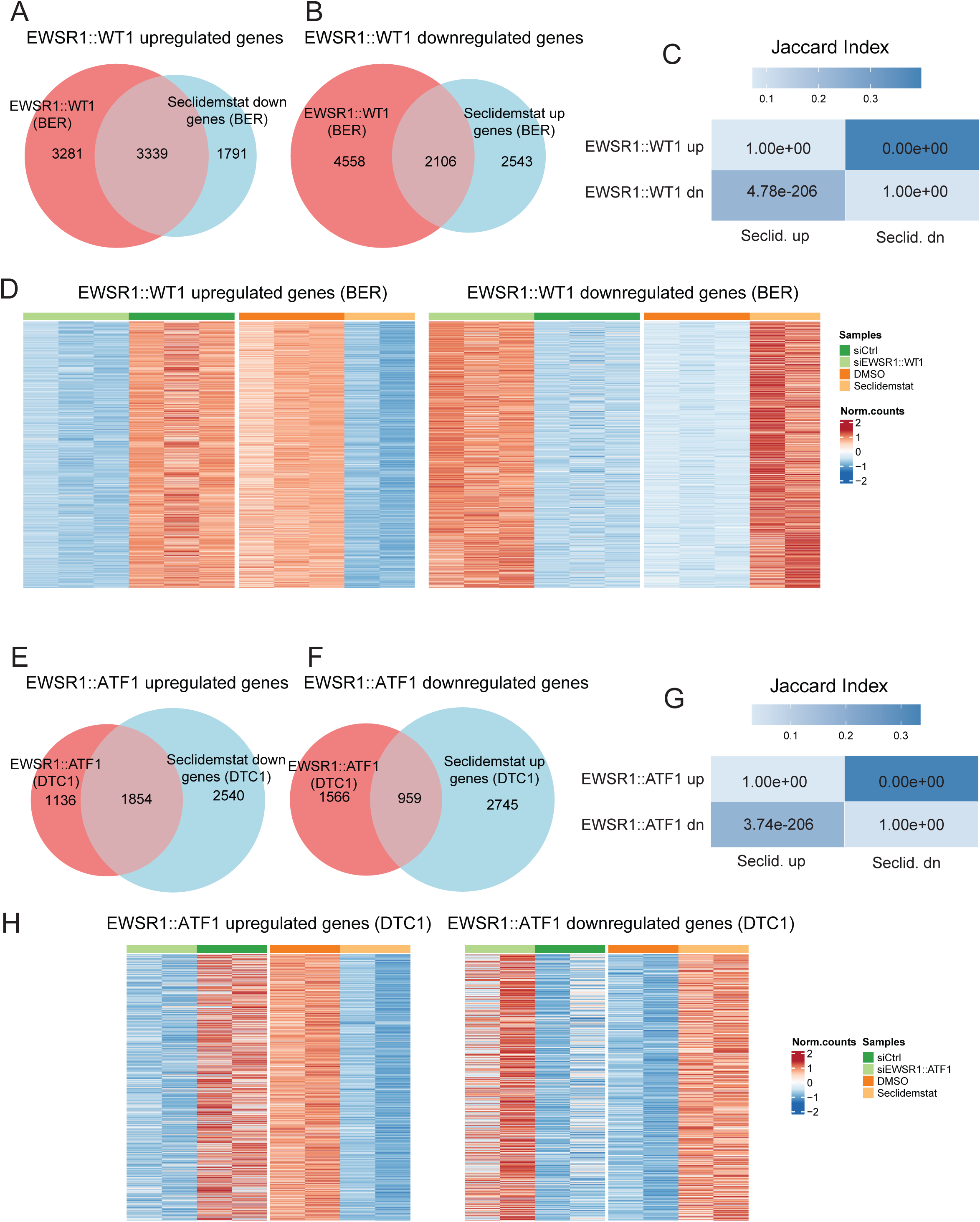
Seclidemstat reverses the transcriptional signature of EWSR1::WT1 and EWSR1::ATF1. (A-C) Venn overlap analysis of (A) seclidemstat downregulated genes and EWSR1::WT1 activated genes and (B) seclidemstat upregulated genes and EWSR1::WT1 repressed genes in BER cells with the Jaccard index and p-values of overlap (numbers inside cell) shown in (C). (D) Heatmap analysis showing the effect of seclidemstat treatment on the EWSR1::WT1 transcriptional signature. Each row represents a differentially expressed gene (adjusted p<0.05) and each column is a separate biological replicate. (E-G) Venn overlap analysis of (E) seclidemstat downregulated genes and EWSR1::ATF1 activated genes and (F) seclidemstat upregulated genes and EWSR1::ATF1 repressed genes in DTC1 cells with the Jaccard index and p-values of overlap (numbers inside cell) shown in (G). (H) Heatmap analysis showing the effect of seclidemstat treatment on the EWSR1::ATF1 transcriptional signature. Each row represents a differentially expressed gene (adjusted p<0.05) and each column is a separate biological replicate.

## DISCUSSION

Taken together, we found that seclidemstat shows potent activity against cells derived from patient tumors with multiple different FET fusion oncogenes. Similar to prior reports in both Ewing sarcoma and DSRCT,^29,45^ cell viability was affected by both of the noncompetitive inhibitors, SP-2509 and seclidemstat, but not OG-L002 indicating that for all of the tumor types tested inhibition of the catalytic function of LSD1 is not a sufficient anti-tumor strategy. One possible explanation for this has been demonstrated by Sehrawat, *et. al.*, 2017^46^, where treatment with SP-2509, but no catalytic LSD1 inhibitor, disrupted interactions between LSD1 and a critical co-regulator ZNF217. It is possible that similar anti-scaffolding activity may be important in FET-rearranged cells and such activity disrupts either the LSD1-ZNF217 interaction or another interaction yet to be defined. Others have suggested that SP-2509 is a compound with pan-assay interference inhibition^47^ (PAINS^48,49^) and may have non-specific off-target effects that confer cytotoxic activity. However, SP-2513 contains the same hydrazide core, with a 2-chloro substituted for the 2-hydroxy of SP-2509 and this compound shows no activity in cell-based assays. Alternatively, there may still be off-target effects related to the *N’*-(2-hydroxybenzilidene)hydrazide core that have not been characterized but nonetheless lead to reversal of FET-fusion protein activity.

This is the largest study to date comparing the transcriptional activity of different FET fusion proteins analyzed together. We were interested to see both a significant portion of fusion-mediated gene regulation unique to each cell line tested as well as a reasonably large number of commonly regulated genes across the 5 cell lines tested. This was particularly true for fusion-activated genes, with 847 commonly upregulated gene targets. While the fusions from each disease have different TF DBDs in the C-terminal portion of the protein, they may nonetheless employ common transcriptional regulatory mechanisms. For example, the EWS LCD is thought to mediate a number of its gene regulatory functions^10,25,50,51^, a new genomic approach to probe the regions of the genome that interact with the EWS LCD has revealed other TFs possibly enriched with fusion proteins like EWSR1::FLI1.^52^ This may prove to be true for other FET-fusion proteins. There were fewer common FET-fusion repressed genes. The overlap may reflect commonalities in cells of origins, while the differences may reflect repression of developmental and signaling pathways unique to each cell of origin for each type of disease and within each patient. Indeed, in our pathway analysis we found several classes of developmental genes and signaling pathways to be commonly repressed.

Many pathways were commonly upregulated across fusions. We were particularly interested to see ribosome biogenesis as highly upregulated genes in all fusion types. This has been described in Ewing sarcoma, with other transcriptomic analyses showing elevated expression of a similar pathway^53^, and also in ultrastructural studies finding an abundance of free ribosomes.^54,55^ Abundant ribosomes are also seen in other small round blue cell tumors.^56^ We also found upregulation of RNA processing genes and pathways related to replication, ssDNA binding and helicase activity. These pathways are consistent with changes in the biology of RNA processing as a result of loss of an allele of one of the RNA-binding FET proteins, and the subsequent alterations in RNA processing caused by interaction of the FET LCD with various RNA processing proteins, like RNA helicase A and the splicing machinery.^57–59^ Additionally, the expression of EWSR1::FLI1 has been shown to drive significant replication stress in Ewing sarcoma cells through elevated levels of transcription, genome wide induction of R-loops, and physical sequestration of proteins involved in resolving replication stress.^43^ Whether other FET fusion oncoproteins affect similar biological pathways in other tumor types is yet to be fully elucidated. We also found transcriptional regulators and co-regulators to be upregulated by all of the fusions and this is consistent with other reports that upregulation of transcriptional regulators is critical for EWSR1::FLI1-mediated oncogenic transformation.^36^

In line with our prior findings for Ewing sarcoma^18^, treatment with noncompetitive LSD1 inhibitors reverses the global transcriptional activity of FET fusion proteins. A significant number of genes modulated by treatment with seclidemstat were unique to each cell line, and this is consistent with uniquely regulated genes being the most prevalent gene class when comparing all fusions. However, pathway analysis showed significant functional overlap in the pathways regulated by seclidemstat treatment in all the tested cell lines. Notably, signatures related to proliferation (cell cycle, replication), RNA processing, and stem cell pathways were downregulated by seclidemstat. Cell signaling and various developmental gene classes were upregulated by seclidemstat. Taken together, these results comprise an orthogonal analytical approach supporting the finding that seclidemstat effectively reverses the oncogenic phenotypes induced by FET fusion proteins. Seclidemstat thus showed comparable pharmacodynamic activity to SP-2509 in these preclinical assay systems, with significant disruption of FET fusion protein function and LSD1. Seclidemstat is currently in clinical trials for patients with Ewing sarcoma and related tumors, as well as MDS and CMML. In interim results from NCT0300649, treatment with seclidemstat shows the potential to prolong time to tumor progression for Ewing sarcoma patients in combination with topotecan and cyclophosphamide, particularly in first relapsed patients. Further prospective, randomized controlled studies are needed to confirm this. Based on our findings that seclidemstat blocks the transcriptional function of other FET fusion proteins, seclidemstat treatment may also be beneficial in other FET-rearranged sarcomas. Additional pharmacokinetic and pharmacodynamic studies are needed to understand whether the current seclidemstat dosing strategy used in the clinic affects FET fusion protein activity in patients and how this might be leveraged in the context of other treatment modalities. SP-2509 has been shown to synergize with HDAC inhibitors in Ewing sarcoma^60^ and this may be an interesting combination to explore. In interim data from NCT04734990 researcher report a 50% objective response and 90% probability of 11 month survival in MDS or CMML patients treated with seclidemstat in combination with azacitidine, additionally supporting the combination of these two epigenetic modalities. Additionally, others have found that the process of tumors becoming resistant to chemotherapy confers collateral sensitivity to SP-2509^61^ such that seclidemstat could be incorporated into an evolutionary treatment framework for sarcoma.^62^ Seclidemstat thus remains a promising new treatment strategy for patients with FET-rearranged sarcomas and future studies are required to understand how to best use this compound in the clinic.

## ACKNOWLEDGEMENTS

We thank Ali Snedden, Yuan Shang, and John Burian at the High-Performance Computing group at Nationwide Children’s Hospital for their support; and the Institute for Genomic Medicine at Nationwide Children’s Hospital for sequencing support. We also acknowledge resources from the OSU Comprehensive Cancer Center (OSUCCC) Medicinal Chemistry Shared Resource (MCSR). We are grateful to the members of the Theisen Lab for comments and discussion of this manuscript during its preparation. This research was supported by institutional startup funds awarded to E.R.T., research project funds awarded by Salarius Pharmaceuticals to E.R.T., and NIH T32 CA269052 to A.B.

