## Supplementary Figures for "Seclidemstat blocks the transcriptional function of multiple FET-fusion oncoproteins"

#### SUPPLEMENTARY FIGURE LEGENDS

**Supplementary Figure 1.** (A-G) Additional Ewing sarcoma replicate dose response curves for seclidemstat (red/circle), SP-2509 (black/closed square), SP-2513 (blue/triangle), and OG-L002 (gray/open square) in (A) A673, (B,C) SK-N-MC, (D,E) TC32, and (F,G) TTC-466 cells. Each graph displays data for a single biological replicate. Mean values of 3 technical replicates are shown with standard deviation. Calculated curves of best fit are also shown.

**Supplementary Figure 2.** (A-G) Additional DSRCT, clear cell sarcoma, and myxoid liposarcoma replicate dose response curves for seclidemstat (red/circle), SP-2509 (black/closed square), SP-2513 (blue/triangle), and OG-L002 (gray/open square) in (A) JN-DSRCT-1, (B,C) BER, (D) SU-CCS-1, (E,F) DTC1, (G) 1765-92, (H,**I**) 402-91, and (J,K) DL221 cells. Each graph displays data for a single biological replicate. Mean values of 3 technical replicates are shown with standard deviation. Calculated curves of best fit are also shown.

**Supplementary Figure 3.** (A,B) Visualization of the fusion calls from the EnFusion pipeline analysis of RNA-seq data show (A) *EWSR1::FLI1* in A673 cells and (B) *EWSR1::ERG* in TTC-466 cells.

**Supplementary Figure 4.** (A,B) Visualization of the fusion calls from the EnFusion pipeline analysis of RNA-seq data show *EWSR1::WT1* in (A) JN-DSRCT-1 and (B) BER cells.

**Supplementary Figure 5.** (A,B) Visualization of the fusion calls from the EnFusion pipeline analysis of RNA-seq data show *EWSR1::ATF1* in (A) SU-CCS-1 and (B) DTC1 cells.

**Supplementary Figure 6.** (A-C) Visualization of the fusion calls from the EnFusion pipeline analysis of RNA-seq data show *FUS::DDIT3* in (A) 1765-92, (B) 402-91, and (C) DL221 cells.

**Supplementary Figure 7.** UMAP plot showing clustering of all analyzed RNA-seq samples.

**Supplementary Figure 8.** (A-C) Venn overlap analysis of SP-2509 and seclidemstat (A) up- and (B) downregulated genes in A673 cells with the Jaccard index and p-values of overlap shown in (C). (D,E) Gene set enrichment analysis of (D) SP-2509 downregulated and (E) SP-2509 upregulated genes with seclidemstat gene regulation as the rank-ordered list. Normalized enrichment score (NES), p-value, and multiple hypothesis adjusted p-values are shown in inset tables.

**Supplementary Figure 9.** (A-C) Venn overlap analysis of (A) EWSR1::FLI1 activated, LSD1 activated, SP-2509 downregulated, and seclidemstat downregulated genes; and (B) EWSR1::FLI1 repressed, LSD1 repressed, SP-2509 upregulated, and seclidemstat upregulated genes in A673 cells with the Jaccard index and p-values of overlap shown in (C). (D-G) Gene set enrichment analysis of (D) LSD1 activated, (E) LSD1 repressed, (F) EWSR1::FLI1 activated and (G) EWSR1::FLI1 repressed genes with seclidemstat gene regulation as the rank-ordered list. Normalized enrichment score (NES), p-value, and multiple hypothesis adjusted p-values are shown in inset tables.

**Supplementary Figure 10.** Pathway analysis for EWSR1::FLI1, LSD1, SP-2509 and seclidemstat regulated genes visualized with a dot plot using MSigDB curated gene signatures.

**Supplementary Figure 11.** (A,B) Pathway analysis for seclidemstat regulated genes in all 9 cell lines tested visualized with a dot plot using (A) gene ontology biological process and (B) gene ontology molecular function gene signatures.

**Supplementary Figure 12.** (A) Western blot showing depletion of endogenous EWSR1::ERG protein in TTC-466 cells and rescue with either empty vector, 3X-FLAG-tagged EWSR1::ERG, or 3X-FLAG-tagged EWSR1::FLI1 in cells used for downstream agar and RNA-seq assays. Whole cell lysates were used. The 3X-FLAG tag causes a slight upward shift in the observed molecular weight of the rescue constructs. (B,C)

(B) Quantification of soft agar assays with representative agar replicates shown in (C). (D) Principal component analysis of gene expression in cells with either endogenous EWSR1::ERG (iLuc+empty vector), EWSR1::ERG1 depletion (iERG+empty vector), or rescue of EWSR1::ERG depletion with either 3X-FLAG EWSR1::ERG (iERG+EWSR1::ERG) or 3X-FLAG EWSR1::FL1 (iERG+EWSR1::FLI1). Principal component 2 on the y-axis is plotted against principal component 1 on the x-axis. Different cell conditions are depicted in different colors and different replicates are represented with different shapes.

**Supplementary Figure 13.** (A,B) Pathway analysis for FET fusion regulated genes in all 5 cell lines tested visualized with a dot plot using (A) gene ontology biological process and (B) gene ontology molecular function gene signatures.

**Supplementary Figure 5.** (A-C) Venn overlap analysis of (A) seclidemstat downregulated genes and EWSR1::WT1 activated genes and (B) seclidemstat upregulated genes and EWSR1::WT1 repressed genes in JN-DSRCT-1 cells with the Jaccard index and p-values of overlap shown in (C). (D) Heatmap analysis showing the effect of seclidemstat treatment on the EWSR1::WT1 transcriptional signature. Each row represents a differentially expressed gene (adjusted $p<0.05$) and each column is a separate biological replicate. (E-G) Venn overlap analysis of (E) seclidemstat downregulated genes and EWSR1::ERG activated genes and (F) seclidemstat upregulated genes and EWSR1::ERG repressed genes in TTC-466 cells with the Jaccard index and p-values of overlap shown in (G). (H) Heatmap analysis showing the effect of seclidemstat treatment on the EWSR1::ERG1 transcriptional signature. Each row represents a differentially expressed gene (adjusted $p<0.05$) and each column is a separate biological replicate.

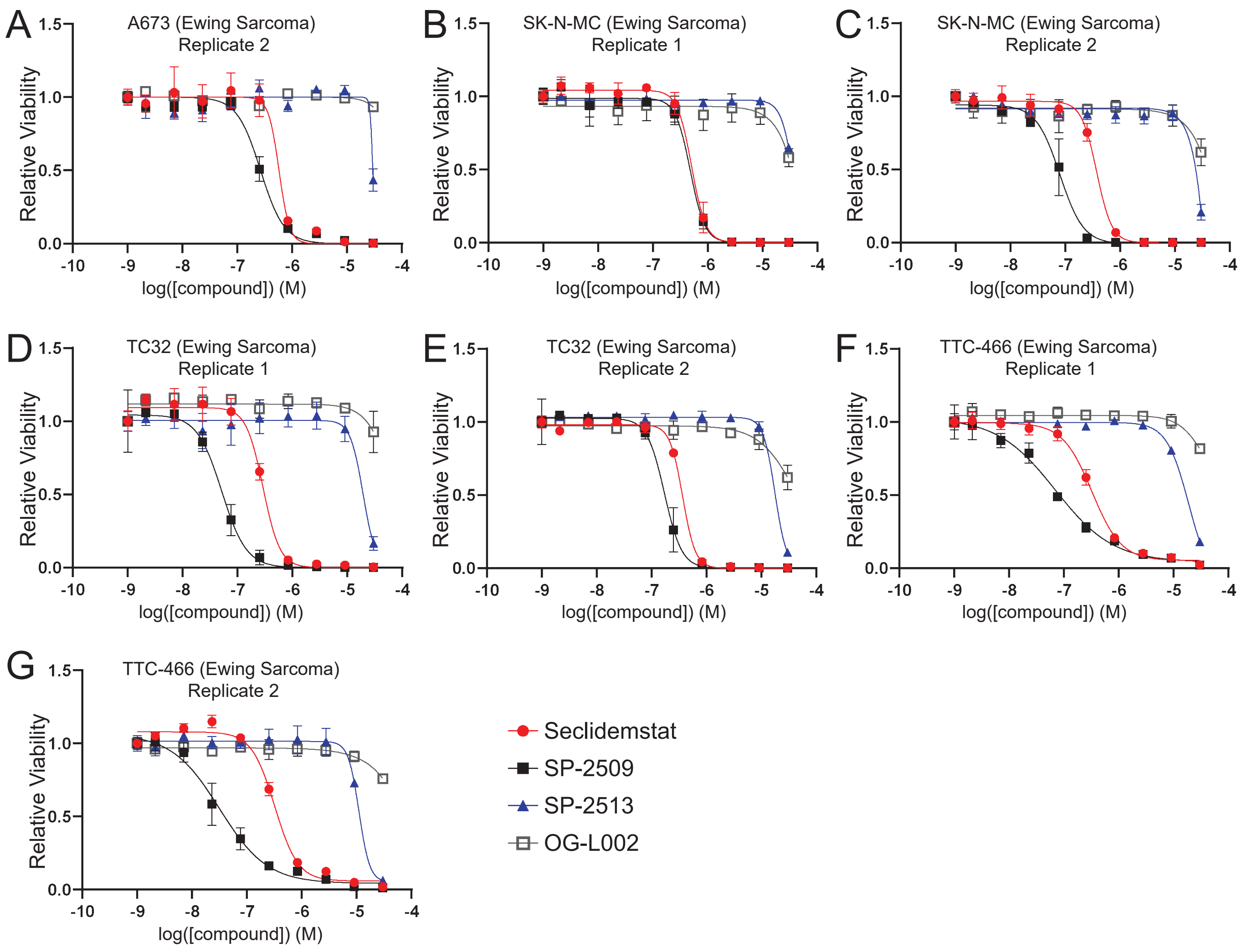
A
A673 (Ewing Sarcoma)
Replicate 2
B
SK-N-MC (Ewing Sarcoma)
Replicate 1
C
SK-N-MC (Ewing Sarcoma)
Replicate 2
D
TC32 (Ewing Sarcoma)
Replicate 1
E
TC32 (Ewing Sarcoma)
Replicate 2
F
TTC-466 (Ewing Sarcoma)
Replicate 1
G
TTC-466 (Ewing Sarcoma)
Replicate 2
Relative Viability
log([compound]) (M)
Seclidemstat
SP-2509
SP-2513
OG-L002

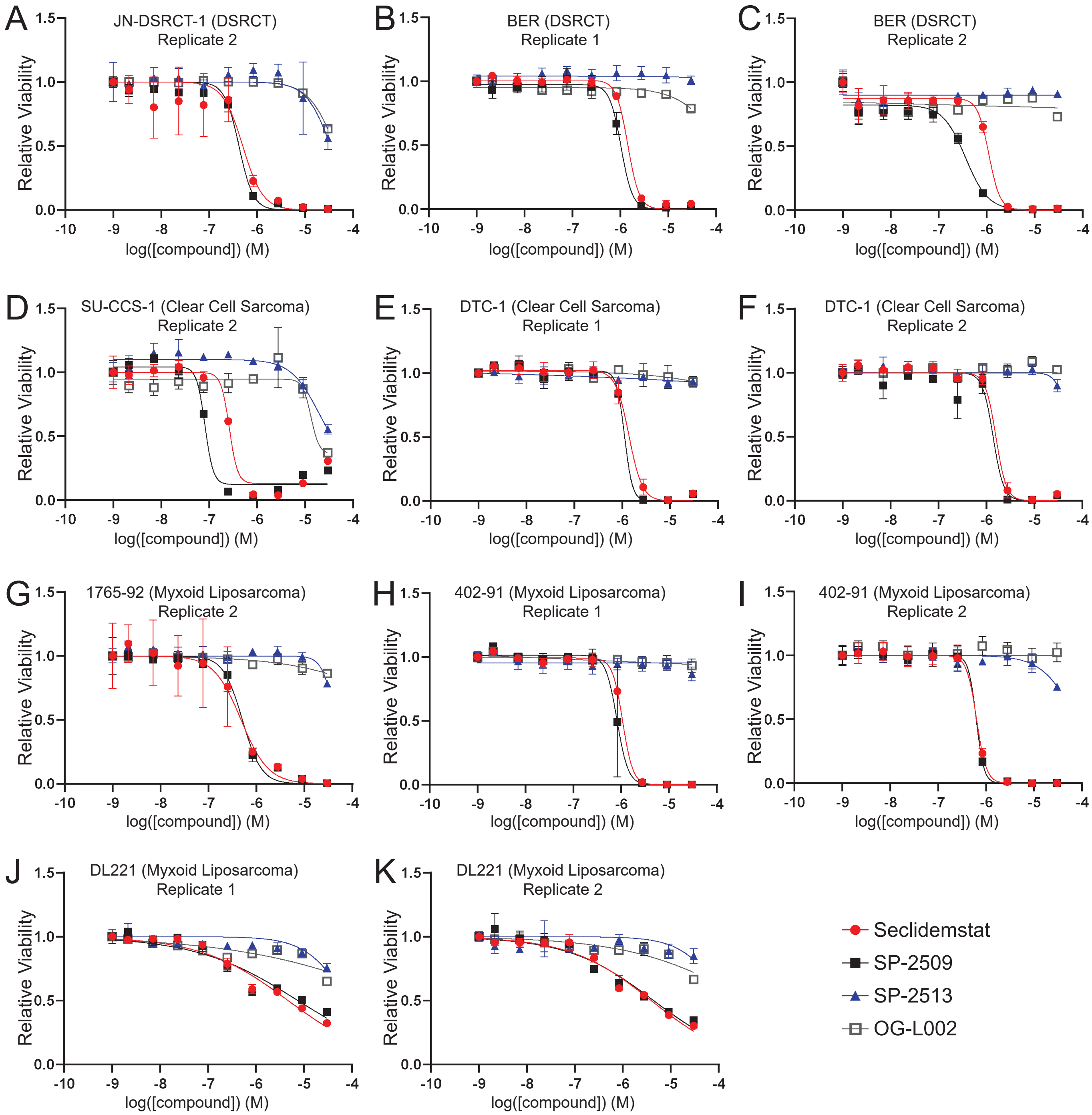

A
JN-DSRCT-1 (DSRCT)
Replicate 2
B
BER (DSRCT)
Replicate 1
C
BER (DSRCT)
Replicate 2
D
SU-CCS-1 (Clear Cell Sarcoma)
Replicate 2
E
DTC-1 (Clear Cell Sarcoma)
Replicate 1
F
DTC-1 (Clear Cell Sarcoma)
Replicate 2
G
1765-92 (Myxoid Liposarcoma)
Replicate 2
H
402-91 (Myxoid Liposarcoma)
Replicate 1
I
402-91 (Myxoid Liposarcoma)
Replicate 2
J
DL221 (Myxoid Liposarcoma)
Replicate 1
K
DL221 (Myxoid Liposarcoma)
Replicate 2
Relative Viability
log([compound]) (M)
Seclidemstat
SP-2509
SP-2513
OG-L002

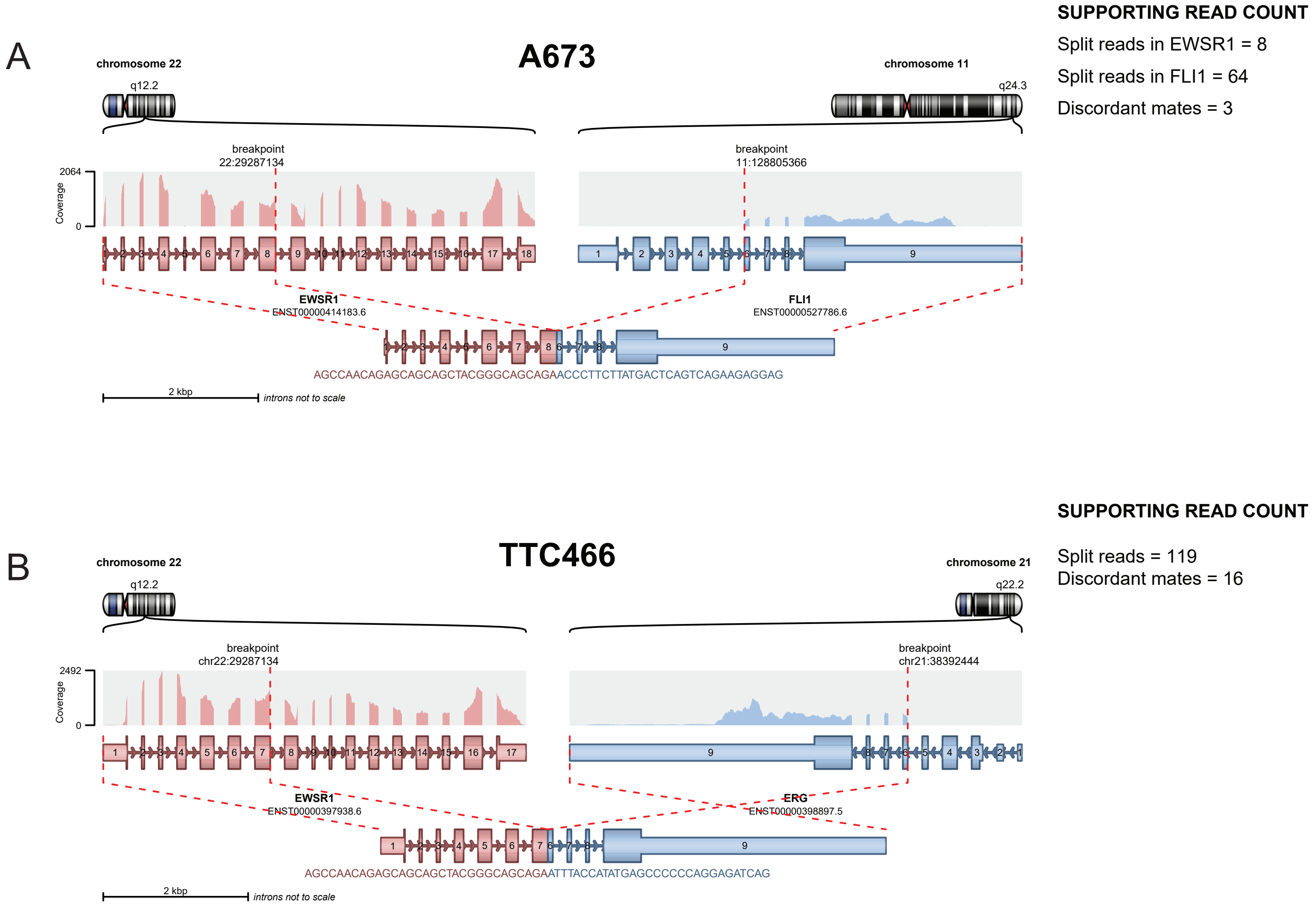
A
A673
chromosome 22
q12.2
chromosome 11
q24.3
SUPPORTING READ COUNT
Split reads in EWSR1 = 8
Split reads in FLI1 = 64
Discordant mates = 3
breakpoint
22:29287134
breakpoint
11:128805366
2064
Coverage
0
EWSR1
ENST00000414183.6
FLI1
ENST00000527786.6
AGCCAACAGAGCAGCAGCTACGGGCAGCAGAACCCTTCTTATGACTCAGTCAGAAGAGGAG
2 kbp
introns not to scale
B
TTC466
chromosome 22
q12.2
chromosome 21
q22.2
SUPPORTING READ COUNT
Split reads = 119
Discordant mates = 16
breakpoint
chr22:29287134
breakpoint
chr21:38392444
2492
Coverage
0
EWSR1
ENST00000397938.6
ERG
ENST00000398897.5
AGCCAACAGAGCAGCAGCTACGGGCAGCAGAATTTACCATATGAGCCCCCCAGGAGATCAG
2 kbp
introns not to scale

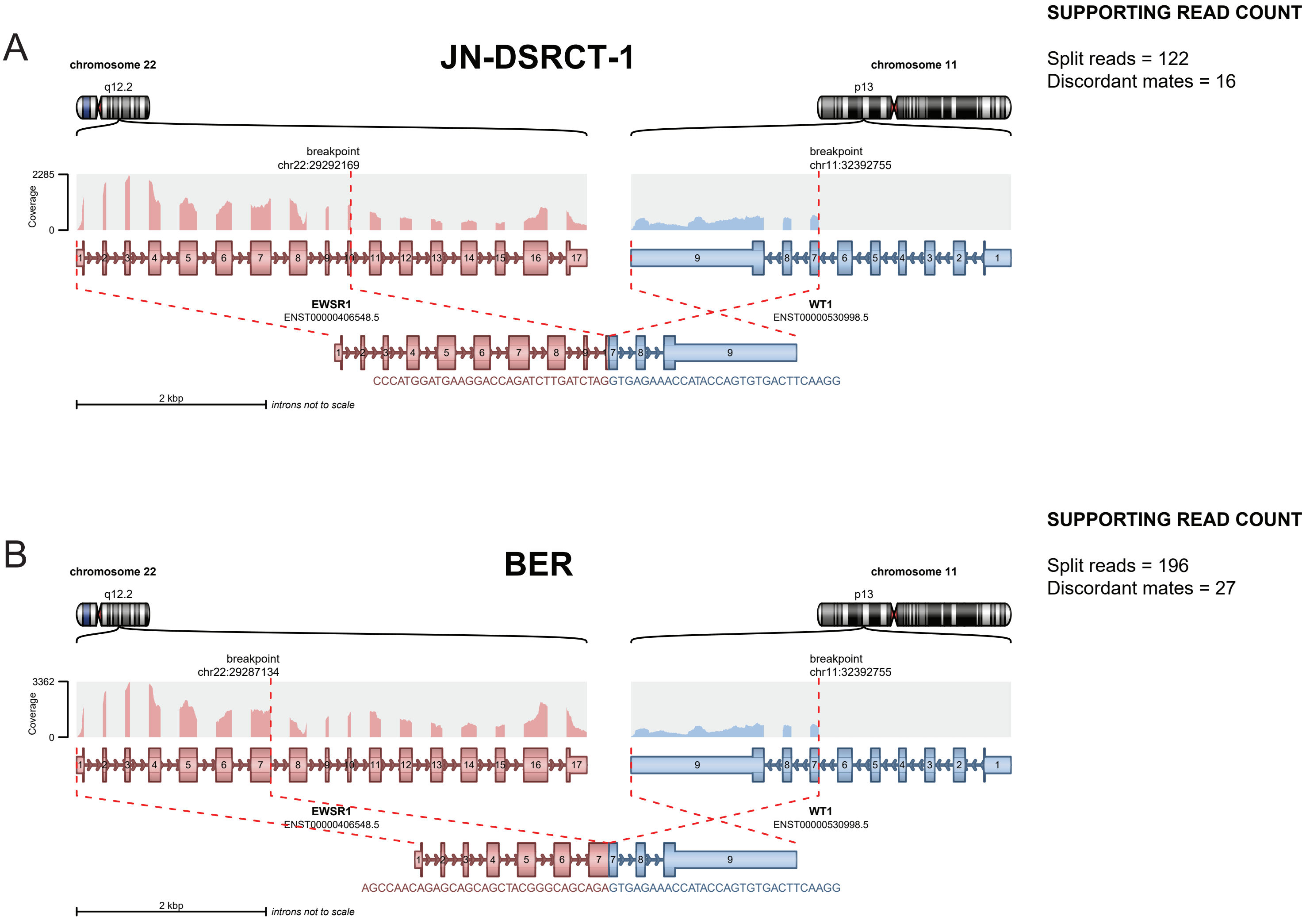
A
JN-DSRCT-1
SUPPORTING READ COUNT
Split reads = 122
Discordant mates = 16
chromosome 22
q12.2
chromosome 11
p13
breakpoint
chr22:29292169
breakpoint
chr11:32392755
2285
Coverage
0
EWSR1
ENST00000406548.5
WT1
ENST00000530998.5
CCCATGGATGAAGGACCAGATCTTGATCTAGGTGAGAAACCATACCAGTGTGACTTCAAGG
2 kbp
introns not to scale
B
BER
SUPPORTING READ COUNT
Split reads = 196
Discordant mates = 27
chromosome 22
q12.2
chromosome 11
p13
breakpoint
chr22:29287134
breakpoint
chr11:32392755
3362
Coverage
0
EWSR1
ENST00000406548.5
WT1
ENST00000530998.5
AGCCAACAGAGCAGCAGCTACGGGCAGCAGAGTGAGAAACCATACCAGTGTGACTTCAAGG
2 kbp
introns not to scale

A

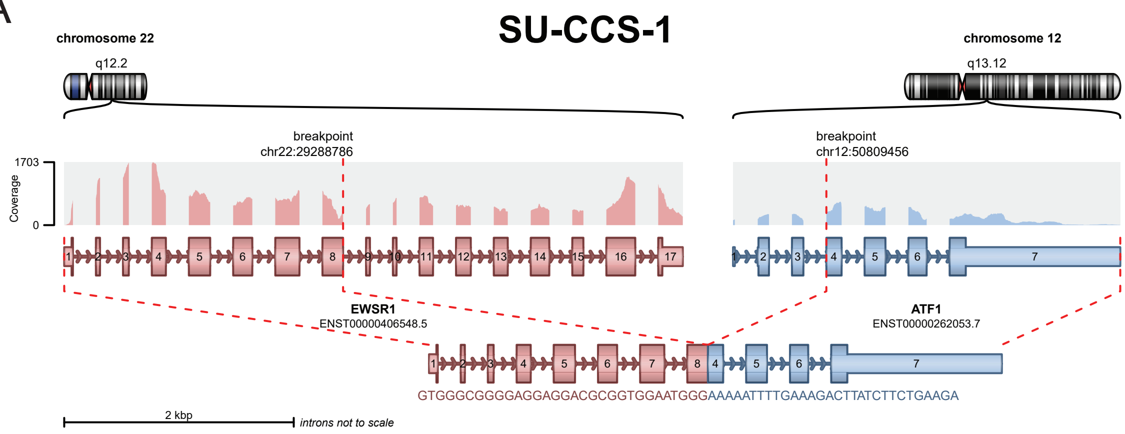


**SUPPORTING READ COUNT**

Split reads = 66
Discordant mates = 1

B

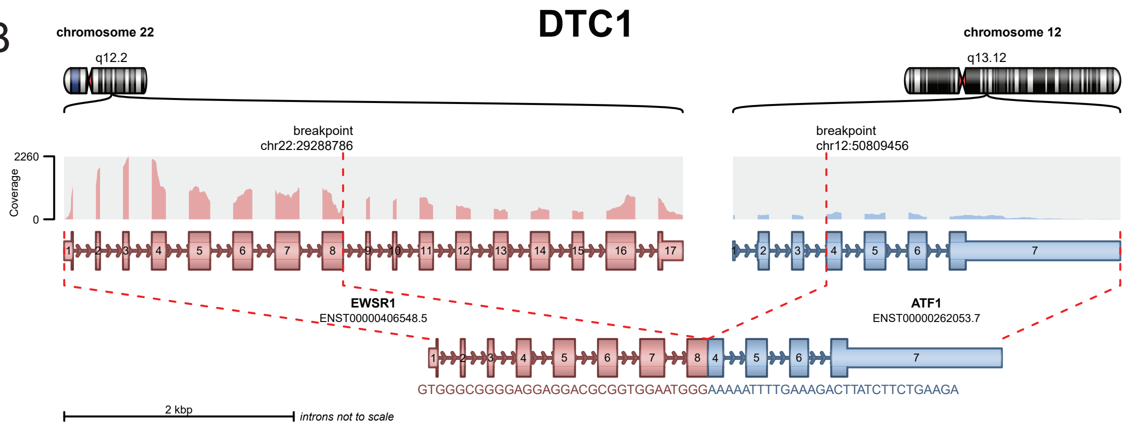


**SUPPORTING READ COUNT**

Split reads = 17
Discordant mates = 3

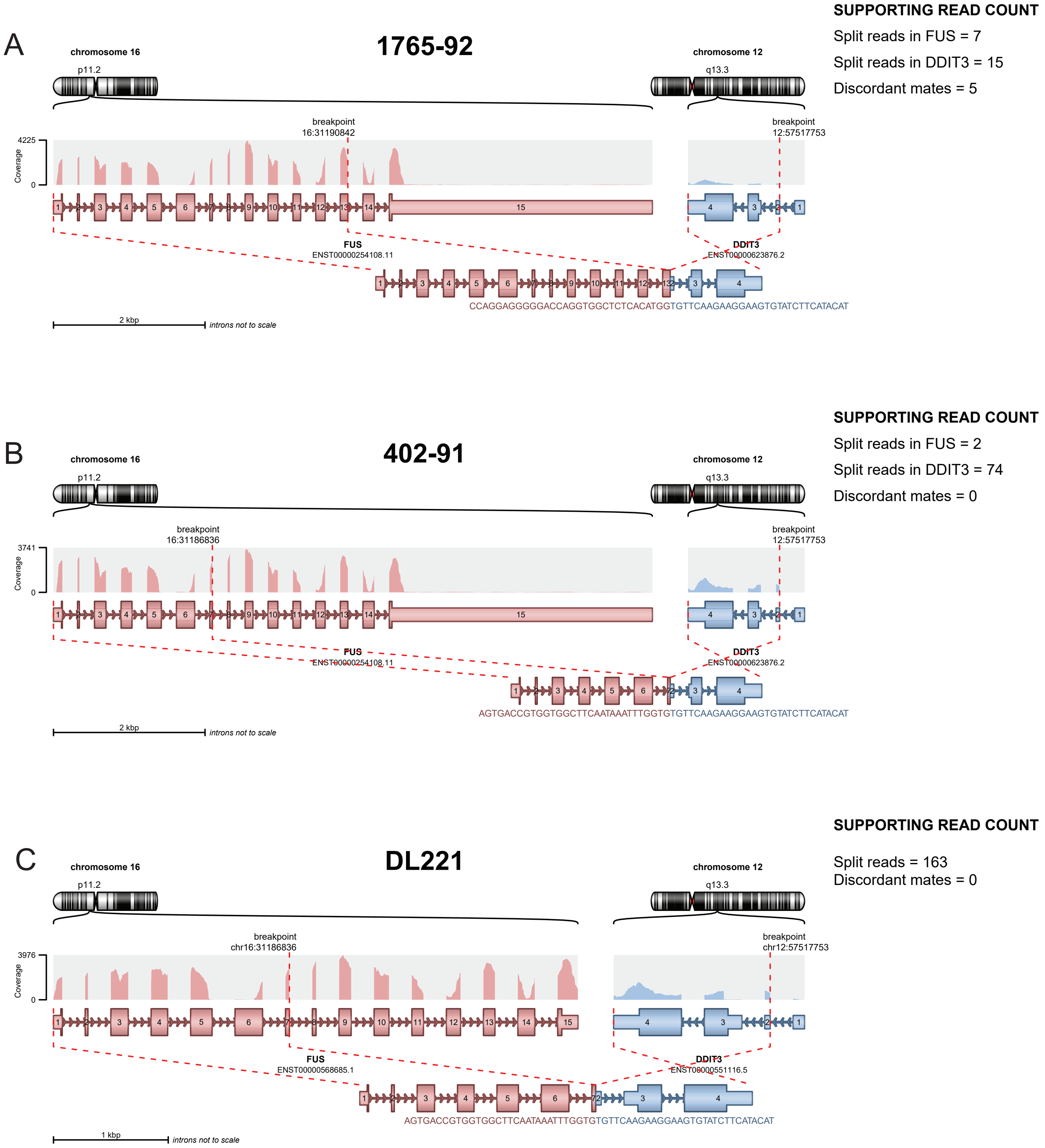

A
1765-92
SUPPORTING READ COUNT
Split reads in FUS = 7
Split reads in DDIT3 = 15
Discordant mates = 5
chromosome 16
p11.2
chromosome 12
q13.3
breakpoint
16:31190842
breakpoint
12:57517753
4225
Coverage
0
FUS
ENST00000254108.11
DDIT3
ENST00000623876.2
CCAGGAGGGGGACCAGGTGGCTCTCACATGGTGTTCAAGAAGGAAGTGTATCTTCATACAT
2 kbp
introns not to scale
B
402-91
SUPPORTING READ COUNT
Split reads in FUS = 2
Split reads in DDIT3 = 74
Discordant mates = 0
chromosome 16
p11.2
chromosome 12
q13.3
breakpoint
16:31186836
breakpoint
12:57517753
3741
Coverage
0
FUS
ENST00000254108.11
DDIT3
ENST00000623876.2
AGTGACCGTGGTGGCTTCAATAAATTTGGTGTGTTCAAGAAGGAAGTGTATCTTCATACAT
2 kbp
introns not to scale
C
DL221
SUPPORTING READ COUNT
Split reads = 163
Discordant mates = 0
chromosome 16
p11.2
chromosome 12
q13.3
breakpoint
chr16:31186836
breakpoint
chr12:57517753
3976
Coverage
0
FUS
ENST00000568685.1
DDIT3
ENST00000551116.5
AGTGACCGTGGTGGCTTCAATAAATTTGGTGTGTTCAAGAAGGAAGTGTATCTTCATACAT
1 kbp
introns not to scale

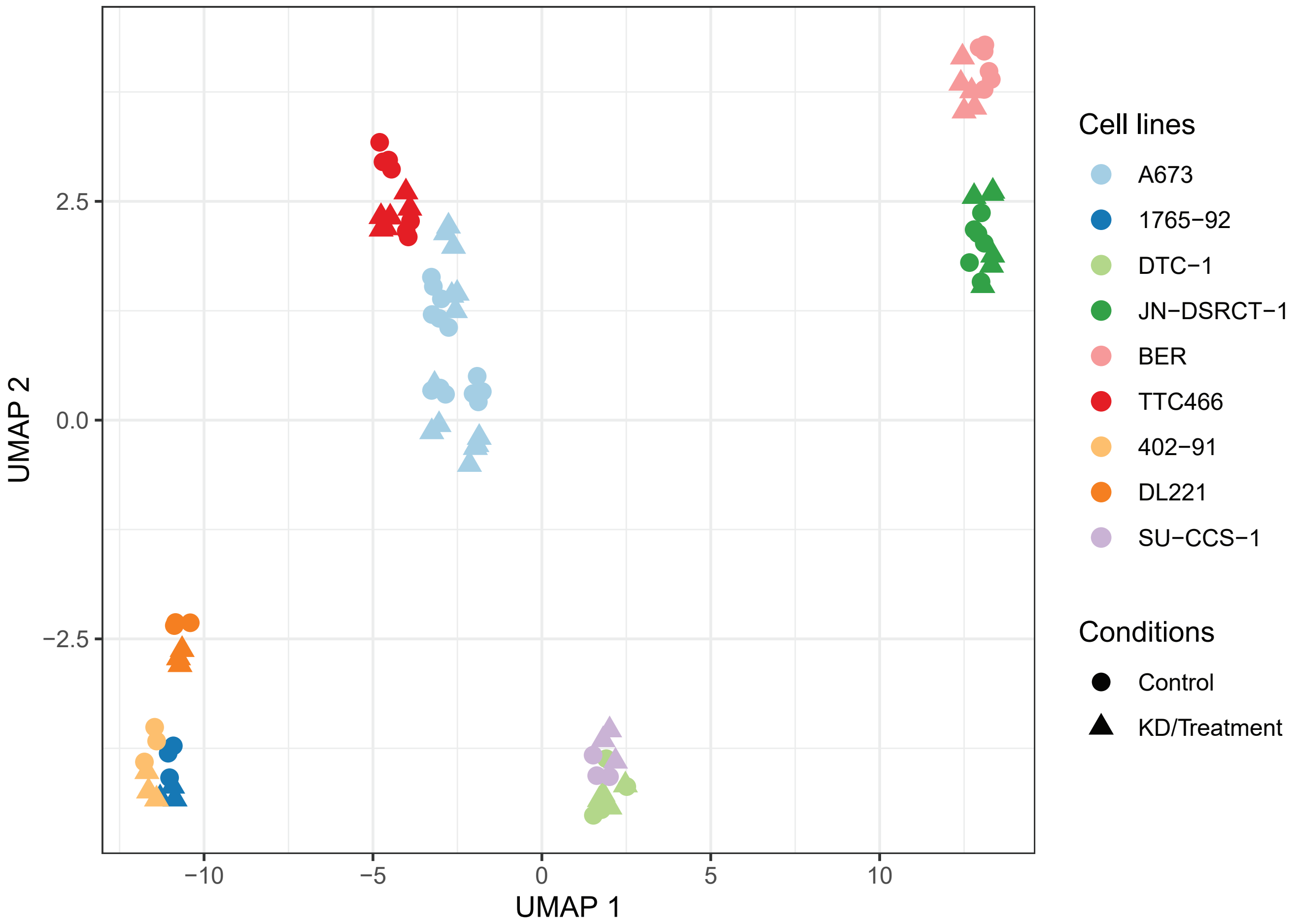
Cell lines
A673
1765−92
DTC−1
JN−DSRCT−1
BER
TTC466
402−91
DL221
SU−CCS−1
Conditions
Control
KD/Treatment
UMAP 1
UMAP 2
−10
−5
0
5
10
2.5
0.0
−2.5

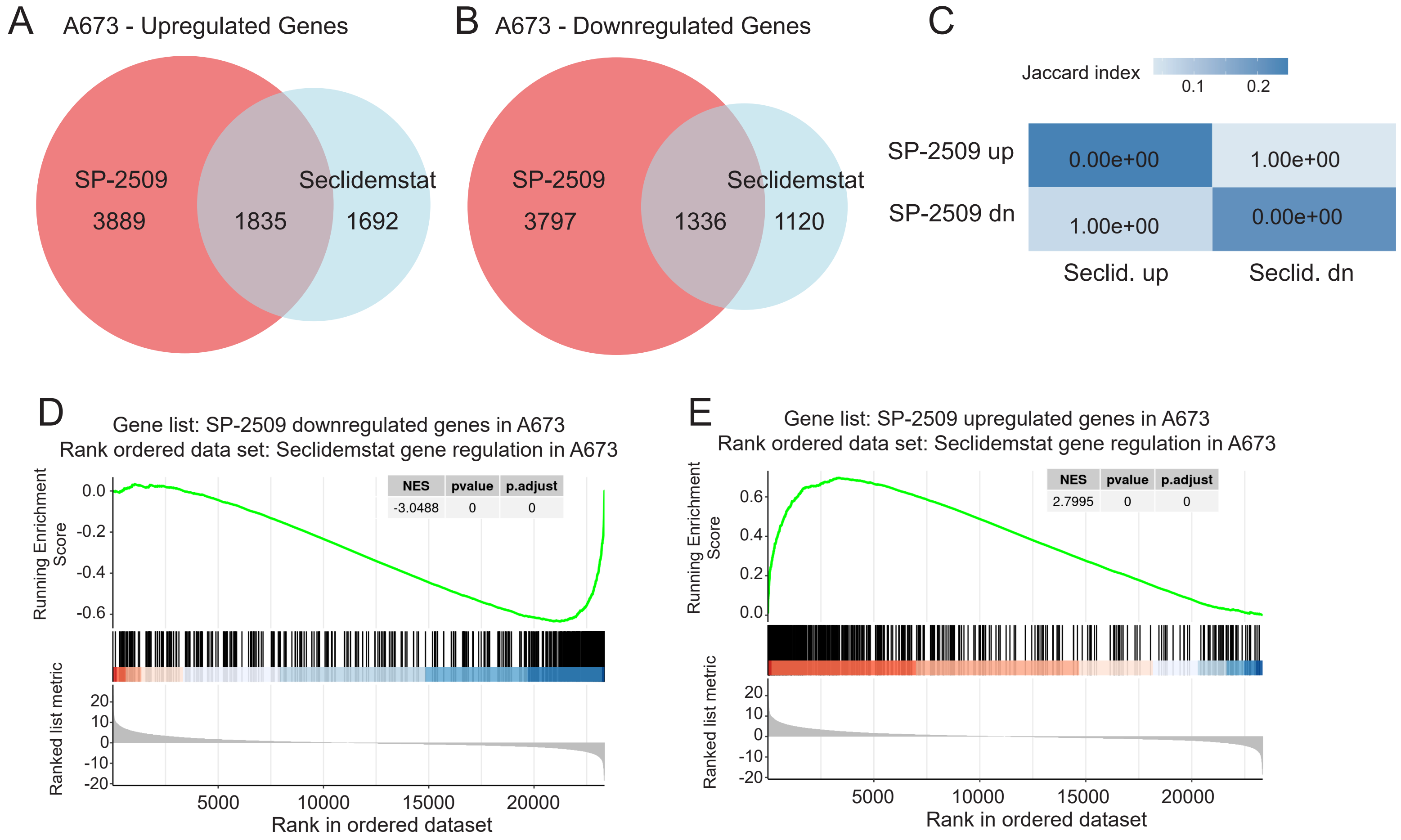
A
A673 - Upregulated Genes
SP-2509
3889
1835
Seclidemstat
1692
B
A673 - Downregulated Genes
SP-2509
3797
1336
Seclidemstat
1120
C
Jaccard index
0.1
0.2
SP-2509 up
SP-2509 dn
0.00e+00
1.00e+00
1.00e+00
0.00e+00
Seclid. up
Seclid. dn
D
Gene list: SP-2509 downregulated genes in A673
Rank ordered data set: Seclidemstat gene regulation in A673
NES
pvalue
p.adjust
-3.0488
0
0
Running Enrichment Score
Ranked list metric
Rank in ordered dataset
E
Gene list: SP-2509 upregulated genes in A673
Rank ordered data set: Seclidemstat gene regulation in A673
NES
pvalue
p.adjust
2.7995
0
0
Running Enrichment Score
Ranked list metric
Rank in ordered dataset

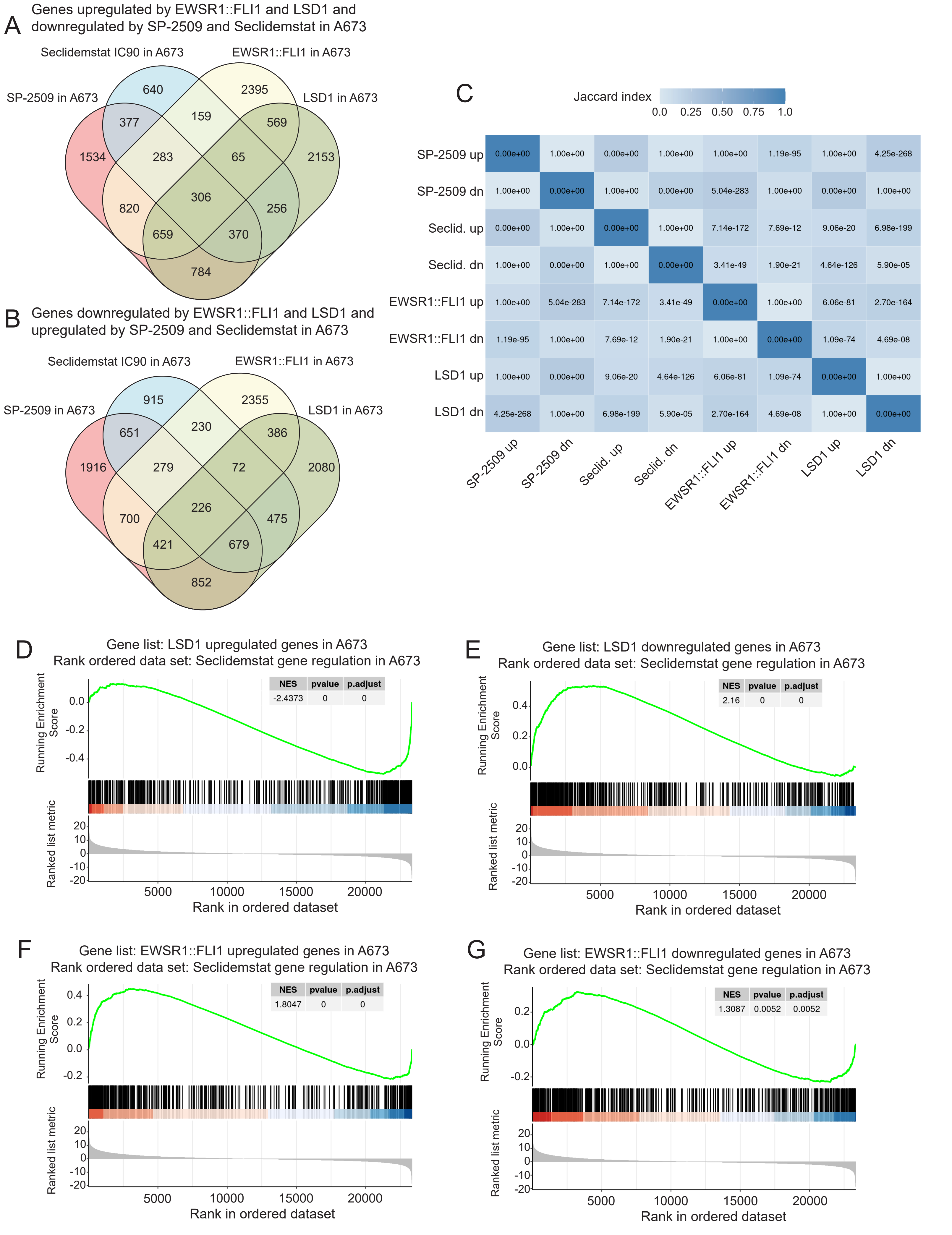
A
Genes upregulated by EWSR1::FLI1 and LSD1 and downregulated by SP-2509 and Seclidemstat in A673
Seclidemstat IC90 in A673
EWSR1::FLI1 in A673
SP-2509 in A673
LSD1 in A673
640
2395
377
159
569
1534
283
65
2153
306
820
256
659
370
784
B
Genes downregulated by EWSR1::FLI1 and LSD1 and upregulated by SP-2509 and Seclidemstat in A673
Seclidemstat IC90 in A673
EWSR1::FLI1 in A673
SP-2509 in A673
LSD1 in A673
915
2355
651
230
386
1916
279
72
2080
226
700
475
421
679
852
C
Jaccard index
0.0 0.25 0.50 0.75 1.0
SP-2509 up
SP-2509 dn
Seclid. up
Seclid. dn
EWSR1::FLI1 up
EWSR1::FLI1 dn
LSD1 up
LSD1 dn
0.00e+00 1.00e+00 0.00e+00 1.00e+00 1.00e+00 1.19e-95 1.00e+00 4.25e-268
1.00e+00 0.00e+00 1.00e+00 0.00e+00 5.04e-283 1.00e+00 0.00e+00 1.00e+00
0.00e+00 1.00e+00 0.00e+00 1.00e+00 7.14e-172 7.69e-12 9.06e-20 6.98e-199
1.00e+00 0.00e+00 1.00e+00 0.00e+00 3.41e-49 1.90e-21 4.64e-126 5.90e-05
1.00e+00 5.04e-283 7.14e-172 3.41e-49 0.00e+00 1.00e+00 6.06e-81 2.70e-164
1.19e-95 1.00e+00 7.69e-12 1.90e-21 1.00e+00 0.00e+00 1.09e-74 4.69e-08
1.00e+00 0.00e+00 9.06e-20 4.64e-126 6.06e-81 1.09e-74 0.00e+00 1.00e+00
4.25e-268 1.00e+00 6.98e-199 5.90e-05 2.70e-164 4.69e-08 1.00e+00 0.00e+00
D
Gene list: LSD1 upregulated genes in A673
Rank ordered data set: Seclidemstat gene regulation in A673
NES pvalue p.adjust
-2.4373 0 0
Running Enrichment Score
Ranked list metric
Rank in ordered dataset
E
Gene list: LSD1 downregulated genes in A673
Rank ordered data set: Seclidemstat gene regulation in A673
NES pvalue p.adjust
2.16 0 0
Running Enrichment Score
Ranked list metric
Rank in ordered dataset
F
Gene list: EWSR1::FLI1 upregulated genes in A673
Rank ordered data set: Seclidemstat gene regulation in A673
NES pvalue p.adjust
1.8047 0 0
Running Enrichment Score
Ranked list metric
Rank in ordered dataset
G
Gene list: EWSR1::FLI1 downregulated genes in A673
Rank ordered data set: Seclidemstat gene regulation in A673
NES pvalue p.adjust
1.3087 0.0052 0.0052
Running Enrichment Score
Ranked list metric
Rank in ordered dataset

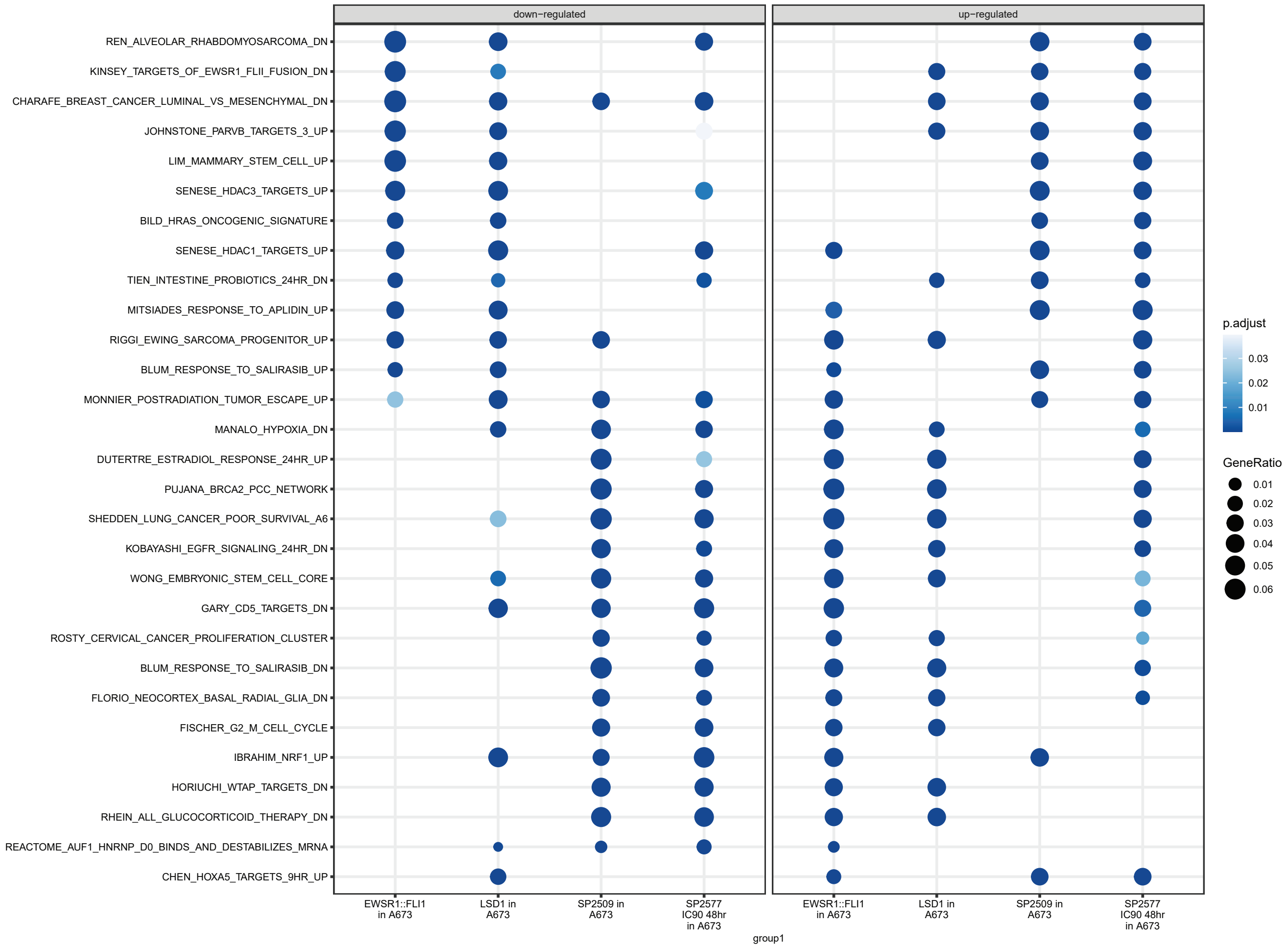
down−regulated
up−regulated
REN_ALVEOLAR_RHABDOMYOSARCOMA_DN
KINSEY_TARGETS_OF_EWSR1_FLII_FUSION_DN
CHARAFE_BREAST_CANCER_LUMINAL_VS_MESENCHYMAL_DN
JOHNSTONE_PARVB_TARGETS_3_UP
LIM_MAMMARY_STEM_CELL_UP
SENESE_HDAC3_TARGETS_UP
BILD_HRAS_ONCOGENIC_SIGNATURE
SENESE_HDAC1_TARGETS_UP
TIEN_INTESTINE_PROBIOTICS_24HR_DN
MITSIADES_RESPONSE_TO_APLIDIN_UP
RIGGI_EWING_SARCOMA_PROGENITOR_UP
BLUM_RESPONSE_TO_SALIRASIB_UP
MONNIER_POSTRADIATION_TUMOR_ESCAPE_UP
MANALO_HYPOXIA_DN
DUTERTRE_ESTRADIOL_RESPONSE_24HR_UP
PUJANA_BRCA2_PCC_NETWORK
SHEDDEN_LUNG_CANCER_POOR_SURVIVAL_A6
KOBAYASHI_EGFR_SIGNALING_24HR_DN
WONG_EMBRYONIC_STEM_CELL_CORE
GARY_CD5_TARGETS_DN
ROSTY_CERVICAL_CANCER_PROLIFERATION_CLUSTER
BLUM_RESPONSE_TO_SALIRASIB_DN
FLORIO_NEOCORTEX_BASAL_RADIAL_GLIA_DN
FISCHER_G2_M_CELL_CYCLE
IBRAHIM_NRF1_UP
HORIUCHI_WTAP_TARGETS_DN
RHEIN_ALL_GLUCOCORTICOID_THERAPY_DN
REACTOME_AUF1_HNRNP_D0_BINDS_AND_DESTABILIZES_MRNA
CHEN_HOXA5_TARGETS_9HR_UP
EWSR1::FLI1 in A673
LSD1 in A673
SP2509 in A673
SP2577 IC90 48hr in A673
EWSR1::FLI1 in A673
LSD1 in A673
SP2509 in A673
SP2577 IC90 48hr in A673
group1
p.adjust
0.03
0.02
0.01
GeneRatio
0.01
0.02
0.03
0.04
0.05
0.06

A

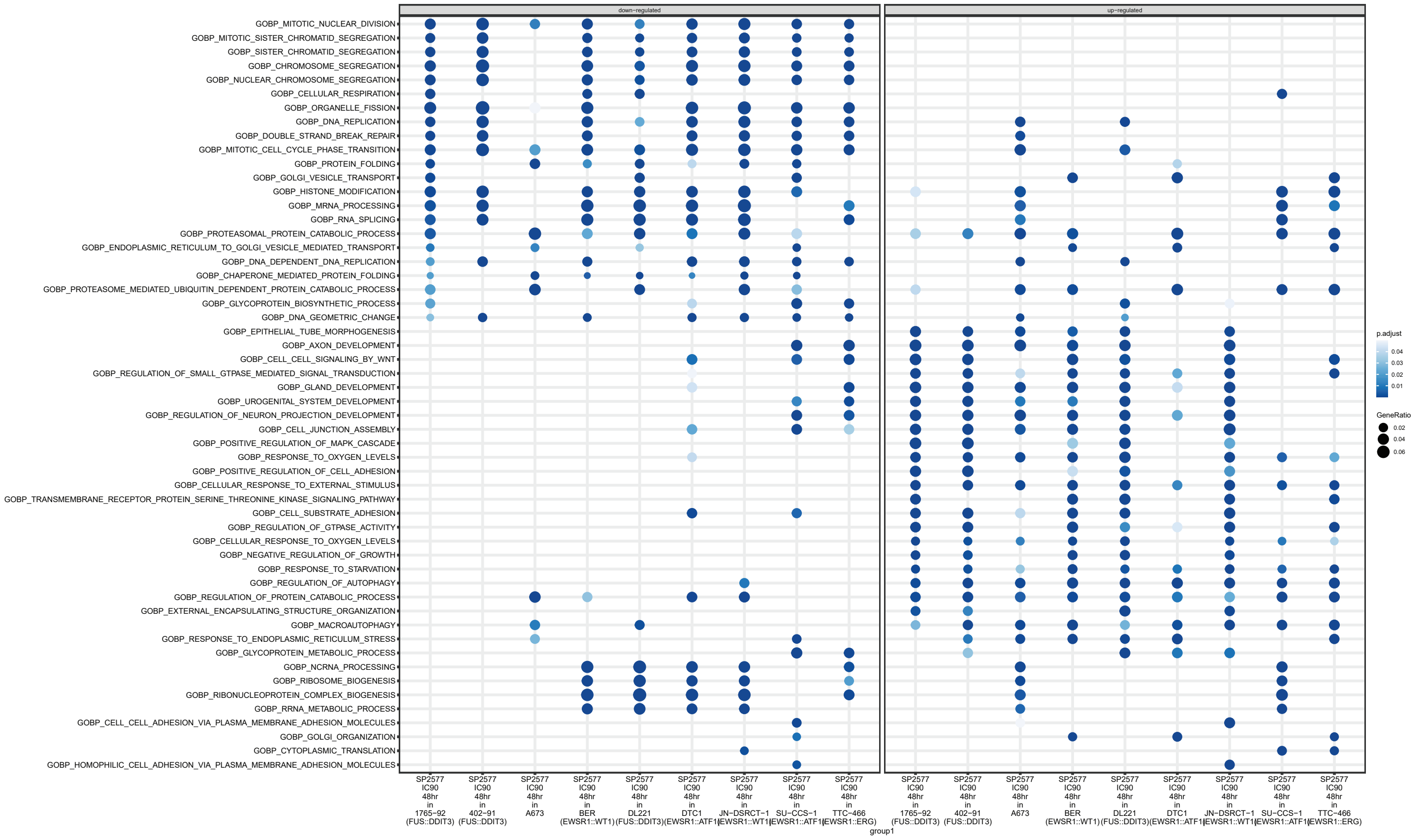
down−regulated
up−regulated
GOBP_MITOTIC_NUCLEAR_DIVISION
GOBP_MITOTIC_SISTER_CHROMATID_SEGREGATION
GOBP_SISTER_CHROMATID_SEGREGATION
GOBP_CHROMOSOME_SEGREGATION
GOBP_NUCLEAR_CHROMOSOME_SEGREGATION
GOBP_CELLULAR_RESPIRATION
GOBP_ORGANELLE_FISSION
GOBP_DNA_REPLICATION
GOBP_DOUBLE_STRAND_BREAK_REPAIR
GOBP_MITOTIC_CELL_CYCLE_PHASE_TRANSITION
GOBP_PROTEIN_FOLDING
GOBP_GOLGI_VESICLE_TRANSPORT
GOBP_HISTONE_MODIFICATION
GOBP_MRNA_PROCESSING
GOBP_RNA_SPLICING
GOBP_PROTEASOMAL_PROTEIN_CATABOLIC_PROCESS
GOBP_ENDOPLASMIC_RETICULUM_TO_GOLGI_VESICLE_MEDIATED_TRANSPORT
GOBP_DNA_DEPENDENT_DNA_REPLICATION
GOBP_CHAPERONE_MEDIATED_PROTEIN_FOLDING
GOBP_PROTEASOME_MEDIATED_UBIQUITIN_DEPENDENT_PROTEIN_CATABOLIC_PROCESS
GOBP_GLYCOPROTEIN_BIOSYNTHETIC_PROCESS
GOBP_DNA_GEOMETRIC_CHANGE
GOBP_EPITHELIAL_TUBE_MORPHOGENESIS
GOBP_AXON_DEVELOPMENT
GOBP_CELL_CELL_SIGNALING_BY_WNT
GOBP_REGULATION_OF_SMALL_GTPASE_MEDIATED_SIGNAL_TRANSDUCTION
GOBP_GLAND_DEVELOPMENT
GOBP_UROGENITAL_SYSTEM_DEVELOPMENT
GOBP_REGULATION_OF_NEURON_PROJECTION_DEVELOPMENT
GOBP_CELL_JUNCTION_ASSEMBLY
GOBP_POSITIVE_REGULATION_OF_MAPK_CASCADE
GOBP_RESPONSE_TO_OXYGEN_LEVELS
GOBP_POSITIVE_REGULATION_OF_CELL_ADHESION
GOBP_CELLULAR_RESPONSE_TO_EXTERNAL_STIMULUS
GOBP_TRANSMEMBRANE_RECEPTOR_PROTEIN_SERINE_THREONINE_KINASE_SIGNALING_PATHWAY
GOBP_CELL_SUBSTRATE_ADHESION
GOBP_REGULATION_OF_GTPASE_ACTIVITY
GOBP_CELLULAR_RESPONSE_TO_OXYGEN_LEVELS
GOBP_NEGATIVE_REGULATION_OF_GROWTH
GOBP_RESPONSE_TO_STARVATION
GOBP_REGULATION_OF_AUTOPHAGY
GOBP_REGULATION_OF_PROTEIN_CATABOLIC_PROCESS
GOBP_EXTERNAL_ENCAPSULATING_STRUCTURE_ORGANIZATION
GOBP_MACROAUTOPHAGY
GOBP_RESPONSE_TO_ENDOPLASMIC_RETICULUM_STRESS
GOBP_GLYCOPROTEIN_METABOLIC_PROCESS
GOBP_NCRNA_PROCESSING
GOBP_RIBOSOME_BIOGENESIS
GOBP_RIBONUCLEOPROTEIN_COMPLEX_BIOGENESIS
GOBP_RRNA_METABOLIC_PROCESS
GOBP_CELL_CELL_ADHESION_VIA_PLASMA_MEMBRANE_ADHESION_MOLECULES
GOBP_GOLGI_ORGANIZATION
GOBP_CYTOPLASMIC_TRANSLATION
GOBP_HOMOPHILIC_CELL_ADHESION_VIA_PLASMA_MEMBRANE_ADHESION_MOLECULES
SP2577 IC90 48hr in 1765−92 (FUS::DDIT3)
SP2577 IC90 48hr in 402−91 (FUS::DDIT3)
SP2577 IC90 48hr in A673
SP2577 IC90 48hr in BER (EWSR1::WT1)
SP2577 IC90 48hr in DL221 (FUS::DDIT3)
SP2577 IC90 48hr in DTC1 (EWSR1::ATF1)
SP2577 IC90 48hr in JN−DSRCT−1 (EWSR1::WT1)
SP2577 IC90 48hr in SU−CCS−1 (EWSR1::ATF1)
SP2577 IC90 48hr in TTC−466 (EWSR1::ERG)
group1
p.adjust
0.04
0.03
0.02
0.01
GeneRatio
0.02
0.04
0.06

B

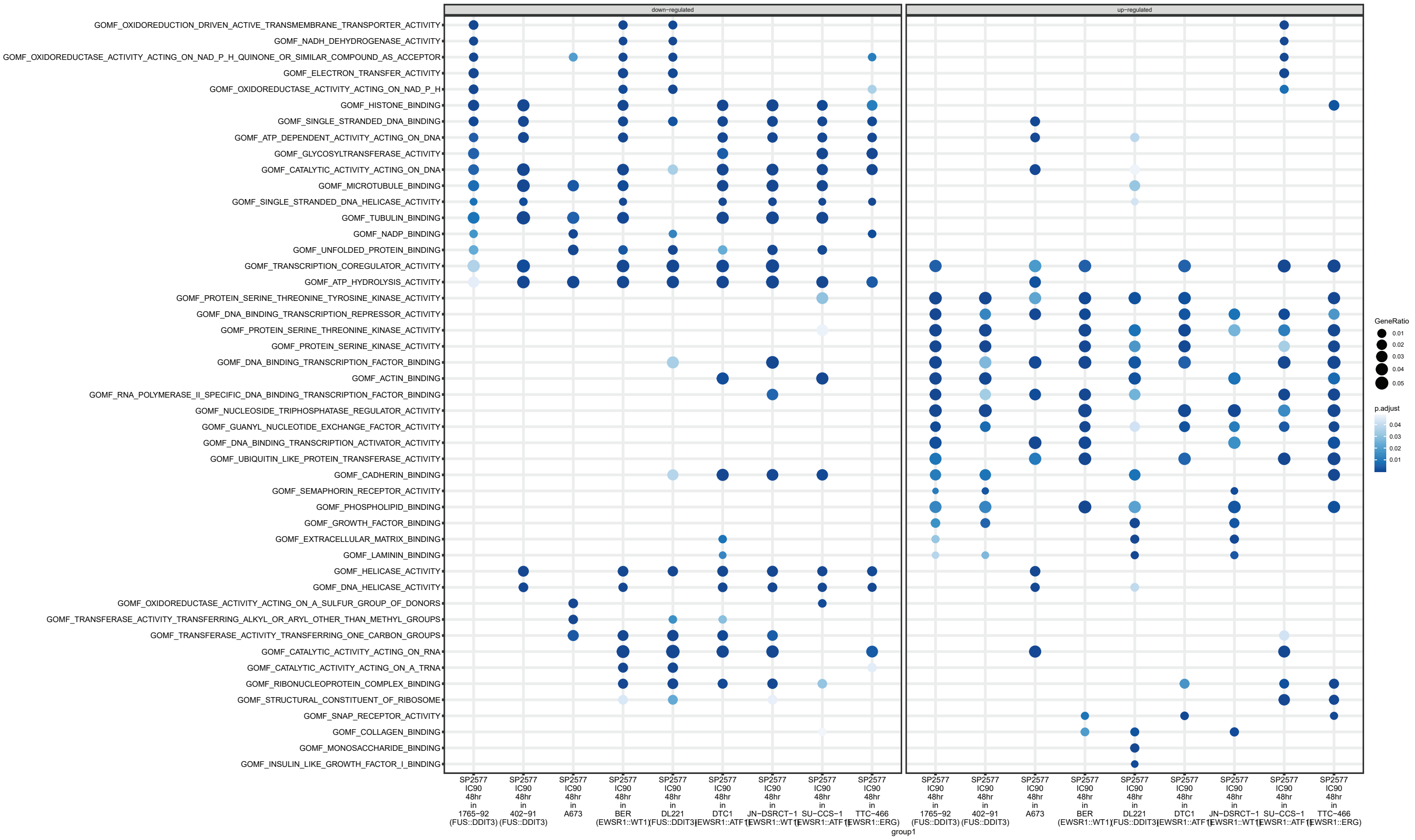
down−regulated
up−regulated
GOMF_OXIDOREDUCTION_DRIVEN_ACTIVE_TRANSMEMBRANE_TRANSPORTER_ACTIVITY
GOMF_NADH_DEHYDROGENASE_ACTIVITY
GOMF_OXIDOREDUCTASE_ACTIVITY_ACTING_ON_NAD_P_H_QUINONE_OR_SIMILAR_COMPOUND_AS_ACCEPTOR
GOMF_ELECTRON_TRANSFER_ACTIVITY
GOMF_OXIDOREDUCTASE_ACTIVITY_ACTING_ON_NAD_P_H
GOMF_HISTONE_BINDING
GOMF_SINGLE_STRANDED_DNA_BINDING
GOMF_ATP_DEPENDENT_ACTIVITY_ACTING_ON_DNA
GOMF_GLYCOSYLTRANSFERASE_ACTIVITY
GOMF_CATALYTIC_ACTIVITY_ACTING_ON_DNA
GOMF_MICROTUBULE_BINDING
GOMF_SINGLE_STRANDED_DNA_HELICASE_ACTIVITY
GOMF_TUBULIN_BINDING
GOMF_NADP_BINDING
GOMF_UNFOLDED_PROTEIN_BINDING
GOMF_TRANSCRIPTION_COREGULATOR_ACTIVITY
GOMF_ATP_HYDROLYSIS_ACTIVITY
GOMF_PROTEIN_SERINE_THREONINE_TYROSINE_KINASE_ACTIVITY
GOMF_DNA_BINDING_TRANSCRIPTION_REPRESSOR_ACTIVITY
GOMF_PROTEIN_SERINE_THREONINE_KINASE_ACTIVITY
GOMF_PROTEIN_SERINE_KINASE_ACTIVITY
GOMF_DNA_BINDING_TRANSCRIPTION_FACTOR_BINDING
GOMF_ACTIN_BINDING
GOMF_RNA_POLYMERASE_II_SPECIFIC_DNA_BINDING_TRANSCRIPTION_FACTOR_BINDING
GOMF_NUCLEOSIDE_TRIPHOSPHATASE_REGULATOR_ACTIVITY
GOMF_GUANYL_NUCLEOTIDE_EXCHANGE_FACTOR_ACTIVITY
GOMF_DNA_BINDING_TRANSCRIPTION_ACTIVATOR_ACTIVITY
GOMF_UBIQUITIN_LIKE_PROTEIN_TRANSFERASE_ACTIVITY
GOMF_CADHERIN_BINDING
GOMF_SEMAPHORIN_RECEPTOR_ACTIVITY
GOMF_PHOSPHOLIPID_BINDING
GOMF_GROWTH_FACTOR_BINDING
GOMF_EXTRACELLULAR_MATRIX_BINDING
GOMF_LAMININ_BINDING
GOMF_HELICASE_ACTIVITY
GOMF_DNA_HELICASE_ACTIVITY
GOMF_OXIDOREDUCTASE_ACTIVITY_ACTING_ON_A_SULFUR_GROUP_OF_DONORS
GOMF_TRANSFERASE_ACTIVITY_TRANSFERRING_ALKYL_OR_ARYL_OTHER_THAN_METHYL_GROUPS
GOMF_TRANSFERASE_ACTIVITY_TRANSFERRING_ONE_CARBON_GROUPS
GOMF_CATALYTIC_ACTIVITY_ACTING_ON_RNA
GOMF_CATALYTIC_ACTIVITY_ACTING_ON_A_TRNA
GOMF_RIBONUCLEOPROTEIN_COMPLEX_BINDING
GOMF_STRUCTURAL_CONSTITUENT_OF_RIBOSOME
GOMF_SNAP_RECEPTOR_ACTIVITY
GOMF_COLLAGEN_BINDING
GOMF_MONOSACCHARIDE_BINDING
GOMF_INSULIN_LIKE_GROWTH_FACTOR_I_BINDING
SP2577 IC90 48hr in 1765−92 (FUS::DDIT3)
SP2577 IC90 48hr in 402−91 (FUS::DDIT3)
SP2577 IC90 48hr in A673
SP2577 IC90 48hr in BER (EWSR1::WT1)
SP2577 IC90 48hr in DL221 (FUS::DDIT3)
SP2577 IC90 48hr in DTC1 (EWSR1::ATF1)
SP2577 IC90 48hr in JN−DSRCT−1 (EWSR1::WT1)
SP2577 IC90 48hr in SU−CCS−1 (EWSR1::ATF1)
SP2577 IC90 48hr in TTC−466 (EWSR1::ERG)
group1
GeneRatio
0.01
0.02
0.03
0.04
0.05
p.adjust
0.04
0.03
0.02
0.01

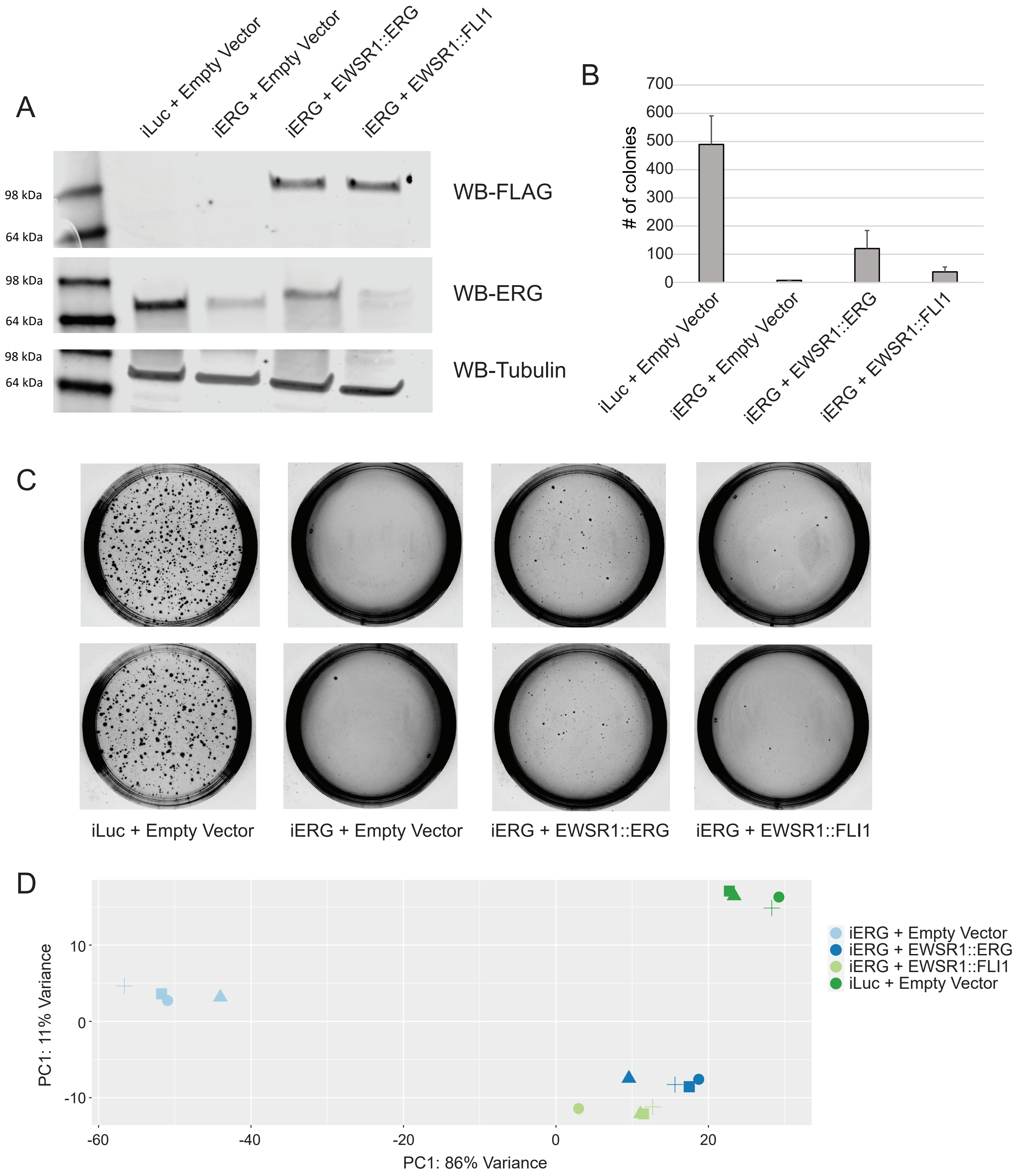

A
iLuc + Empty Vector
iERG + Empty Vector
iERG + EWSR1::ERG
iERG + EWSR1::FLI1
98 kDa
64 kDa
WB-FLAG
98 kDa
64 kDa
WB-ERG
98 kDa
64 kDa
WB-Tubulin
B
### of colonies
700
600
500
400
300
200
100
0
iLuc + Empty Vector
iERG + Empty Vector
iERG + EWSR1::ERG
iERG + EWSR1::FLI1
C
iLuc + Empty Vector
iERG + Empty Vector
iERG + EWSR1::ERG
iERG + EWSR1::FLI1
D
PC1: 11% Variance
10
0
-10
-60
-40
-20
0
20
PC1: 86% Variance
iERG + Empty Vector
iERG + EWSR1::ERG
iERG + EWSR1::FLI1
iLuc + Empty Vector

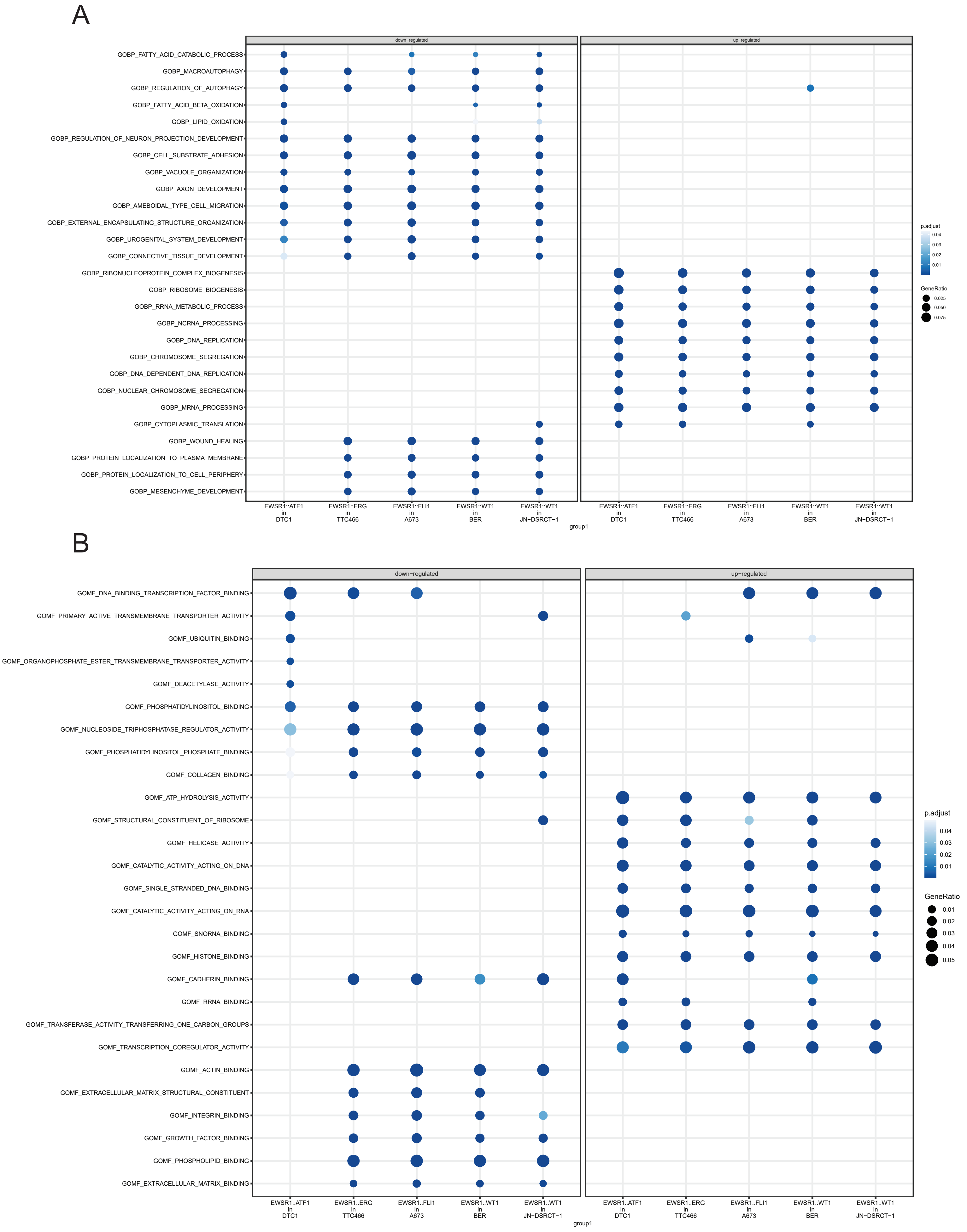
A
down−regulated
up−regulated
GOBP_FATTY_ACID_CATABOLIC_PROCESS
GOBP_MACROAUTOPHAGY
GOBP_REGULATION_OF_AUTOPHAGY
GOBP_FATTY_ACID_BETA_OXIDATION
GOBP_LIPID_OXIDATION
GOBP_REGULATION_OF_NEURON_PROJECTION_DEVELOPMENT
GOBP_CELL_SUBSTRATE_ADHESION
GOBP_VACUOLE_ORGANIZATION
GOBP_AXON_DEVELOPMENT
GOBP_AMEBOIDAL_TYPE_CELL_MIGRATION
GOBP_EXTERNAL_ENCAPSULATING_STRUCTURE_ORGANIZATION
GOBP_UROGENITAL_SYSTEM_DEVELOPMENT
GOBP_CONNECTIVE_TISSUE_DEVELOPMENT
GOBP_RIBONUCLEOPROTEIN_COMPLEX_BIOGENESIS
GOBP_RIBOSOME_BIOGENESIS
GOBP_RRNA_METABOLIC_PROCESS
GOBP_NCRNA_PROCESSING
GOBP_DNA_REPLICATION
GOBP_CHROMOSOME_SEGREGATION
GOBP_DNA_DEPENDENT_DNA_REPLICATION
GOBP_NUCLEAR_CHROMOSOME_SEGREGATION
GOBP_MRNA_PROCESSING
GOBP_CYTOPLASMIC_TRANSLATION
GOBP_WOUND_HEALING
GOBP_PROTEIN_LOCALIZATION_TO_PLASMA_MEMBRANE
GOBP_PROTEIN_LOCALIZATION_TO_CELL_PERIPHERY
GOBP_MESENCHYME_DEVELOPMENT
EWSR1::ATF1 in DTC1
EWSR1::ERG in TTC466
EWSR1::FLI1 in A673
EWSR1::WT1 in BER
EWSR1::WT1 in JN−DSRCT−1
group1
p.adjust
0.04
0.03
0.02
0.01
GeneRatio
0.025
0.050
0.075
B
down−regulated
up−regulated
GOMF_DNA_BINDING_TRANSCRIPTION_FACTOR_BINDING
GOMF_PRIMARY_ACTIVE_TRANSMEMBRANE_TRANSPORTER_ACTIVITY
GOMF_UBIQUITIN_BINDING
GOMF_ORGANOPHOSPHATE_ESTER_TRANSMEMBRANE_TRANSPORTER_ACTIVITY
GOMF_DEACETYLASE_ACTIVITY
GOMF_PHOSPHATIDYLINOSITOL_BINDING
GOMF_NUCLEOSIDE_TRIPHOSPHATASE_REGULATOR_ACTIVITY
GOMF_PHOSPHATIDYLINOSITOL_PHOSPHATE_BINDING
GOMF_COLLAGEN_BINDING
GOMF_ATP_HYDROLYSIS_ACTIVITY
GOMF_STRUCTURAL_CONSTITUENT_OF_RIBOSOME
GOMF_HELICASE_ACTIVITY
GOMF_CATALYTIC_ACTIVITY_ACTING_ON_DNA
GOMF_SINGLE_STRANDED_DNA_BINDING
GOMF_CATALYTIC_ACTIVITY_ACTING_ON_RNA
GOMF_SNORNA_BINDING
GOMF_HISTONE_BINDING
GOMF_CADHERIN_BINDING
GOMF_RRNA_BINDING
GOMF_TRANSFERASE_ACTIVITY_TRANSFERRING_ONE_CARBON_GROUPS
GOMF_TRANSCRIPTION_COREGULATOR_ACTIVITY
GOMF_ACTIN_BINDING
GOMF_EXTRACELLULAR_MATRIX_STRUCTURAL_CONSTITUENT
GOMF_INTEGRIN_BINDING
GOMF_GROWTH_FACTOR_BINDING
GOMF_PHOSPHOLIPID_BINDING
GOMF_EXTRACELLULAR_MATRIX_BINDING
EWSR1::ATF1 in DTC1
EWSR1::ERG in TTC466
EWSR1::FLI1 in A673
EWSR1::WT1 in BER
EWSR1::WT1 in JN−DSRCT−1
group1
p.adjust
0.04
0.03
0.02
0.01
GeneRatio
0.01
0.02
0.03
0.04
0.05

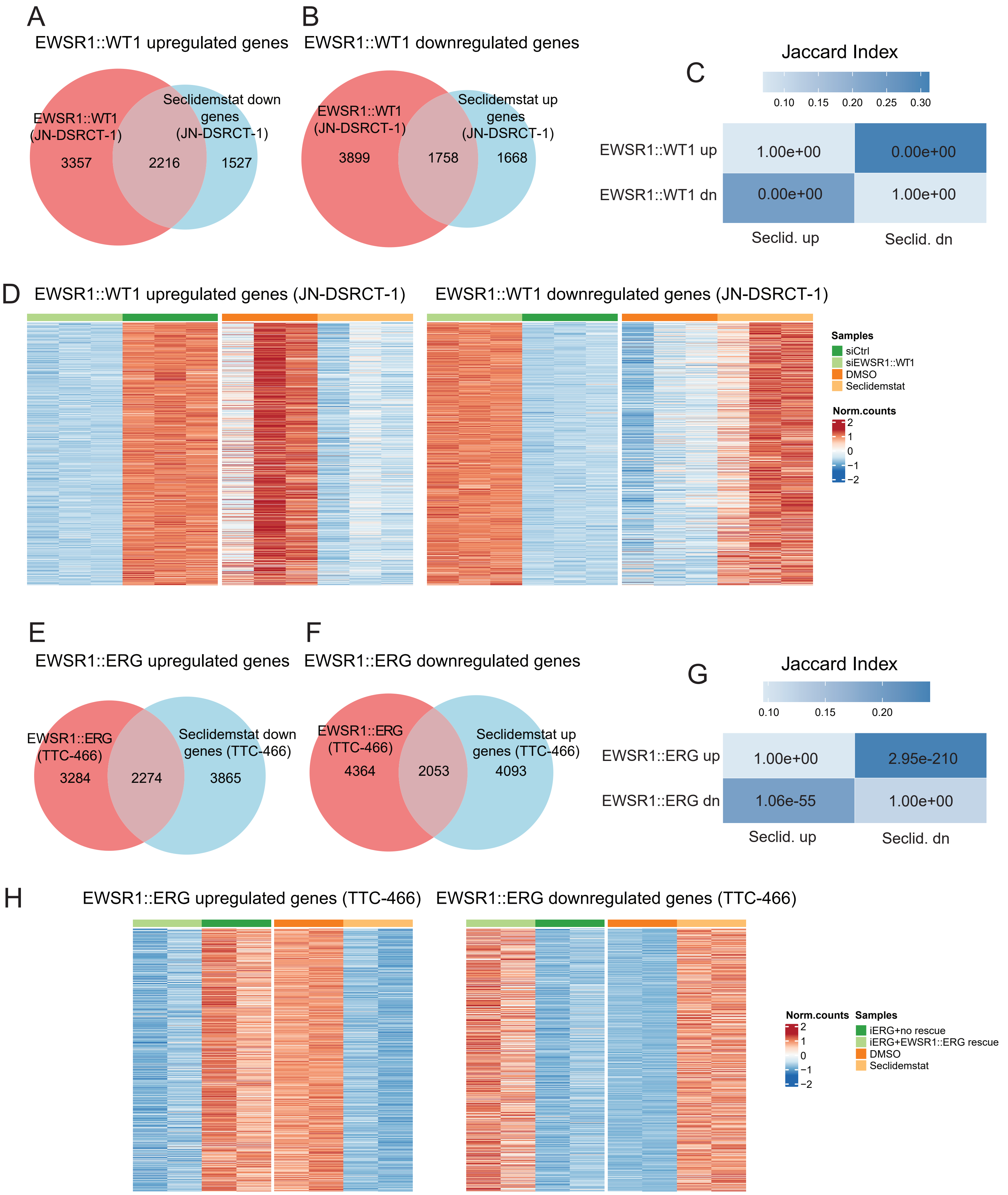
A
EWSR1::WT1 upregulated genes
EWSR1::WT1 (JN-DSRCT-1)
Seclidemstat down genes (JN-DSRCT-1)
3357
2216
1527
B
EWSR1::WT1 downregulated genes
EWSR1::WT1 (JN-DSRCT-1)
Seclidemstat up genes (JN-DSRCT-1)
3899
1758
1668
C
Jaccard Index
0.10
0.15
0.20
0.25
0.30
EWSR1::WT1 up
EWSR1::WT1 dn
1.00e+00
0.00e+00
0.00e+00
1.00e+00
Seclid. up
Seclid. dn
D
EWSR1::WT1 upregulated genes (JN-DSRCT-1)
EWSR1::WT1 downregulated genes (JN-DSRCT-1)
Samples
siCtrl
siEWSR1::WT1
DMSO
Seclidemstat
Norm.counts
2
1
0
−1
−2
E
EWSR1::ERG upregulated genes
EWSR1::ERG (TTC-466)
Seclidemstat down genes (TTC-466)
3284
2274
3865
F
EWSR1::ERG downregulated genes
EWSR1::ERG (TTC-466)
Seclidemstat up genes (TTC-466)
4364
2053
4093
G
Jaccard Index
0.10
0.15
0.20
EWSR1::ERG up
EWSR1::ERG dn
1.00e+00
2.95e-210
1.06e-55
1.00e+00
Seclid. up
Seclid. dn
H
EWSR1::ERG upregulated genes (TTC-466)
EWSR1::ERG downregulated genes (TTC-466)
Norm.counts
2
1
0
−1
−2
Samples
iERG+no rescue
iERG+EWSR1::ERG rescue
DMSO
Seclidemstat
